# Interplay of Cullin5-HECTD3 ubiquitin ligases regulates stability of CRAF mutant associated with hypertrophy

**DOI:** 10.64898/2026.05.23.727451

**Authors:** Somesh Roy, Soumita Mukherjee, Atin K. Mandal

## Abstract

Proteostatic imbalance caused by CRAF (Raf1) kinase mutation alters MAPK signaling and contributes to RASopathies. CRAF proteostasis is maintained by chaperones, Hsp90 which aids in folding, maturation, and activation; whereas Hsp70 recruits ubiquitin ligase CHIP for proteasomal degradation. However, the proteostasis of disease-associated CRAF mutants remains undefined. Here, we identify HECTD3, a HECT-ubiquitin ligase in promoting degradation of hypertrophic CRAF mutants. Interestingly, increased Hsp90 binding to CRAF plays a determinant factor in degradation via HECTD3. Transient increase in Hsp90-CRAF association upon Hsp90 inhibition promote ubiquitination, whereas prolonged inhibition stabilizes mutant CRAF, indicating temporally distinct chaperone functions. Notably, HECTD3 undergoes proteosomal degradation under stress via Cullin-RING ubiquitin ligase Cul5 in a neddylation-dependent manner. Overexpression of Cul5 thus reduces HECTD3 level resulted in stabilization of CRAF^D486N^, while promoting degradation of wild-type CRAF. Therefore, modulation of the Cul5-HECTD3 axis alters hypertrophic signaling in cardiomyocyte cells. These results establish a hierarchical proteostasis mechanism in which Hsp90 binding primes substrate for HECTD3-mediated ubiquitination, and Cul5-controlled ligase turnover collectively determine mutant CRAF fate.

## Introduction

CRAF kinase is a central component of the mitogen-activated protein kinase (MAPK) signaling cascade and regulates key cellular processes such as proliferation, migration and differentiation through the formation of homodimers (CRAF-CRAF) or heterodimers with BRAF [1–3]. Dysregulation of this pathway resulting from germ-line mutations in CRAF has been implicated in several pathological conditions, including cancer and RASopathies such as Noonan and LEOPARD syndromes, which frequently present with hypertrophic cardiomyopathy (HCM) [4–6]. These mutations in CRAF encompass both gain-of-function variants, such as S257L and L613V, and loss-of-function and kinase-inactive variants such as D486N [6–8]. Despite their opposing effects on kinase activity, these mutants have been reported to promote hypertrophic signaling through enhanced association with BRAF and mostly by activating calcineurin-NFAT pathway independent of MAPK signaling, ultimately driving the transcription of hypertrophic genes [6, 8].

The stability and activity of CRAF within the cellular environment are regulated by a network of molecular chaperones, scaffolding proteins and ubiquitin ligases other than various kinase proteins [9–11]. CRAF is a well-established client of the Hsp90 chaperone machinery and requires continuous interaction with Hsp90 and its co-chaperone Cdc37 to maintain proper folding and phosphorylation at Ser621, a modification essential for its maturation, stability and activity [12, 13]. Inhibition of Hsp90 disrupts this chaperone cycle, leading to reduced Ser621 phosphorylation, enhanced association of CRAF with the chaperone complex, and subsequent ubiquitin ligase-mediated degradation [12, 14]. Previous studies have shown that under basal conditions wild-type CRAF is primarily degraded by the Hsp70-associated U-box ubiquitin ligase CHIP, whereas during Hsp90 inhibition degradation is mediated through the HECT domain-containing ubiquitin ligase HECTD3 [15, 16]. Interestingly, kinase-inactive mutants such as D486A have been reported to exhibit reduced Ser621 phosphorylation together with enhanced interaction with Hsp90, suggesting that certain mutant forms of CRAF may resemble the Hsp90-inhibited state of wild-type CRAF [12, 13]. However, how kinase-inactive or hypertrophy-associated CRAF mutants are regulated by the cellular chaperone-ubiquitin system remains unknown. In particular, it remains unclear whether these mutants utilize similar quality-control mechanisms as wild-type CRAF during chaperone perturbation, or whether distinct regulatory pathways influence their stability [13, 16]. Given the reported increase in Hsp90 association observed in certain inactive CRAF mutants, the relationship between chaperone engagement and ubiquitin ligase-mediated degradation of mutant CRAF proteins requires further exploration.

To address this question, the behavior of the kinase-inactive CRAF mutant, D486N was examined under conditions of Hsp90 perturbation and compared with that of wild-type CRAF, with a focus on chaperone association and degradation dynamics. In this study, a striking similarity in chaperone association was observed between wild-type CRAF under Hsp90-inhibited conditions and mutant forms of CRAF described previously [13, 16]. Whereas inhibition of Hsp90 accelerated the degradation of wild-type CRAF, the kinase-inactive mutant D486N displayed a partial recovery in protein levels following prolonged inhibition, suggesting the involvement of compensatory chaperone-dependent regulatory mechanisms. Further investigation identified the ubiquitin ligase HECTD3, previously implicated in the degradation of wild-type CRAF during Hsp90 inhibition, as a regulator of the stability of specific hypertrophy-associated CRAF mutants. Mechanistic analyses further revealed that HECTD3 itself undergoes destabilization under proteotoxic stress conditions, particularly upon Hsp90 inhibition, through a pathway mediated by the Cullin-RING ligase Cul5, by activation through its neddylation. Notably, neddylation-activated Cul5 promoted stabilization of the hypertrophic mutant in a dose-dependent manner, revealing a feedback regulatory mechanism linking Cul5 activity to mutant CRAF stability. Functional analyses further demonstrated that the interplay between HECTD3 and Cul5 influences the stability of hypertrophic CRAF mutants and modulates hypertrophic signaling in cardiomyocyte-derived cells.

## Materials and Methods

### Cell culture and transfection

Human embryonic kidney 293T (HEK293T) cell line, rat cardiomyocytes (H9c2) cell line were purchased from the National Centre for Cell Science (NCCS), Pune, India, and human ventricular cardiomyocytes (AC16) cell line was cultured in DMEM (gibco) supplemented with 10% FBS (gibco), 150 μg/ml Pen-strep, 250 μg/ml Amphotericin-B, 50 μg/ml Gentamycin, and conditioned with 5% CO2 in a humidified CO2 incubator. Cells were split at 80%-90% confluency for regular maintenance. For transient transfection at 50%-60% confluency, cells were transfected with either PEI (polyethylene imine, Polysciences), or Fugene HD (Promega), somethimes with Lipofectamine 2000 (Invitrogen) in Opti-MEM media (gibco). shRNA transfection was carried out with a separate protocol.

### Immunoblotting & Immunoprecipitation

Cells were first lysed in RIPA buffer-I containing 50 mM Tris-HCl (pH 7.5), 150 mM NaCl, 1% Triton-X 100, 0.1% SDS, and 1 mM PMSF with protease inhibitor cocktail for normal immunoblot (not for immunoprecipitation or sh/sg mediated silencing immunoblots); RIPA buffer-II containing 50 mM Tris-HCl (pH 7.5), 150 mM NaCl, 0.8-1% Triton-X 100, and 1 mM PMSF with protease inhibitor cocktail for immunoprecipitation and RIPA buffer-III containing 50 mM Tris-HCl (pH 7.5), 150 mM NaCl, 2% SDS, and 1 mM PMSF with protease inhibitor cocktail for silencing mediated immunoblots. Cell lysates were cleared at 12,000g for 20 min at 4°C. Proteins were estimated using the BCA protein estimation kit (ThermoFisher) and subsequently separated by SDS-PAGE and transferred onto nitrocellulose membrane (Pall Corporation, NY, or Bio-Rad). For immunoprecipitation the cell lysates were incubated with antibodies in an IP dilution Buffer (50 mM Tris, pH 7.5, 150 mM NaCl, 0.2%Triton-X 100, PMSF) overnight at 4°C with gentle rotation. After incubation, protein A-Sepharose (GE Healthcare) bead or protein G magnetic beads (Invitrogen) pre-equilibrated with IP dilution buffer was added and nutated further for 2 hr at 4°C for immunoprecipitation. The beads were then washed three times with IP dilution buffer. Bound proteins were eluted from beads with SDS sample buffer, vortexed, boiled for 5 minutes, and analyzed by western blotting.The proteins were visualized using an ECL detection buffer, and the image was captured in the Chemidoc MP system (Bio-Rad).

### Ubiquitination assay

Cells were transfected with DNA for 48h. Then treated with proteasome inhibitor MG132 for 4h as indicated time points in figure legend before harvesting the cells. The cells were then lysed with cell lysis buffer (50 mM Tris-HCl, pH 7.5, 150 mM NaCl, 0.8% TritonX-100, 1.5% SDS, 1mM PMSF with protease inhibitor cocktail) with sonication. The lysates were diluted 4 times with IP Dilution Buffer (50 mM Tris-HCl, pH 7.5, 150 mM NaCl, 0.8% TritonX-100) and protein was estimated by the BCA method. Corresponding antibody was added to the lysate, and the immunocomplex was incubated at 4°C overnight with gentle shaking. The next day, beads were added to the cell lysate-antibody mixture and incubated at 4°C for 4 hours. The resin was then washed three times with wash buffer (50 mM Tris-HCl, pH 7.5, 150 mM NaCl, 1% TritonX-100, 0.1% SDS) and boiled with 2X SDS loading buffer for immunoblot analysis.

### Real time quantitative PCR

AC16 cells were maintained in low serum (2% horse serum) media for 3-5 days before transfection. After transfection (6h of transfection), transfected cells were maintained in low serum media (2% horse serum) for 24h and cells were harvested for BNP gene expression. For ANP gene expression, after the initial 24 h of low serum, the cells were further maintained for an additional 48 h in serum-free medium to assess the precise effect of the transfected protein. Scrsh & shHERC2 stable *CRAF^-/-^* HEK293T cells were harvested for HERC2 gene expression level. Total RNA was isolated with TRIzol reagent (Invitrogen, 15596018). cDNA was synthesized by using a ABScript II cDNA First Strand Synthesis Kit (Abclonal, RK20400). The RT-qPCR was performed by using Genious 2X SYBR Green Fast qPCR Mix (Abclonal, RK21204) using a Thermocycler Detection System QuantStudio5 (Applied Biosystems, Thermo).

### Confocal microscopy

HEK293T cells were transfected with either PX459 empty vector (i) or sgCul5-1 (ii). Then transfected cells were selected in puromycin (0.75µg/ml) added complete DMEM medium for 10-12 days. Then polyclonal sgCul5-1 HEK293T cells were transfected with pcDNA3-Myc-Cul5^WT^ for 60-72h (iii). For confocal imaging, DNA transfection was carried out at 60% confluence cells and grown on cover slips (without any coating). After 24h of cell seeding on coverslips, cells were fixed with 3.7% formaldehyde, washed with 3 times 1X phosphate-buffered saline (PBS) for 5 min. The cells were then permeabilized with 0.1% triton-X 100 and blocked with 5% BSA for 1h at room temperature (RT). The coverslips washed 2 times with 1X PBS and probed with primary antibody at 4°C overnight. Cells (i & ii) were immunostained with Cul5 antibody (mouse Alexa-594 secondary antibody) and cells (iii) was immunostained with Myc antibody (mouse Alexa-594 secondary antibody). Next day coverslips were treated with mentioned secondary antibody for 1h at RT and then immunostained with Phalloidin labeling probes (Alexa-488) followed by DAPI (Roche). Then cover slips were washed with 3 times 1X PBS and mounted onto glass slide with Vectashield (Vector laboratories, H-1000) and sealed with nail polish. Cells were visualized at 63X oil objective with 4X zoom in a Leica Stellaris-5 confocal microscope.

### Knockout cell line and shRNA mediated stable cell line preparation

CRAF knockout HEK293T cell was generated by CRISPR/Cas9 system, using guide RNA targeting exon 3 of CRAF and cloned into pSpCas9(BB)-2A-Puro (PX459) V2.0, obtained from Addgene (#62988). Cells were then transfected with the cloned plasmid, and the media were changed to complete Dulbecco’s modified Eagle medium after 6 h of transfection. Puromycin (0.75 μg/ml) was added after 48 h of transfection, and the cells were cultured for 12 days before colony picking and monoclonal expansion. Targeted region and predicted off-target exons were checked from monoclone no. 1 (*CRAF^-/-^_Clone-1_*) by sequencing, which is used for the experiments [17].

For the Cul5 knockout polyclonal cell and CRAF-Cul5 double knockout cell, using guide RNA targeting exon 3 of Cul5 (sgCul5-1), and by following the same protocol as for the CRAF knockout cell, monoclone no. 6 (*CRAF^-/-^ Cul5^-/-^*) was checked for experiments.

For the CRAF-Hsp90β double knockout cell, using guide RNA targeting exon 2 of Hsp90β, and by following the same protocol as for the CRAF-Cul5 double knockout cell, monoclone no. 2 (*CRAF^-/-^ Hsp90β^-/-^*) was checked for experiments.

Short hairpin RNA (shRNA) targeted to control and all the ubiquitin ligases was generated by viral particle mediated stable shRNA including scramble and shRNA-pLKO.1-TRC vector by using the shRNA primers.

### Reagents, antibodies, plasmids and primers are given in the supporting information Statistical Analysis

Repeated experiments were statistically analyzed by one-way ANOVA (for more than two groups) or Student’s t test (for two groups). Statistical data was quantified using Prism 9 (GraphPad Software) and the significance of the differences was determined by ± SEM of the quantified values.

## Results

### Hsp90-mediated degradation initiation of CRAF^D486N^

Previous studies have shown that the degradation of wild-type CRAF (CRAF^WT^) is regulated by the molecular chaperone Hsp70 and its co-chaperone Bag1 [15]. However, subsequent work by Noble et al. revealed that this mechanism does not apply to certain inactive CRAF mutants. In particular, the kinase-dead mutant D486A (CRAF^D486A^; mutation in the ‘D’ residue of the DFG motif) does not undergo degradation through Hsp70 or Bag1 [13]. Instead, CRAF^D486A^ displays increased association with Hsp90 compared with CRAF^WT^ [13]. Interestingly, a similar increase in Hsp90 binding has also been observed for CRAF^WT^ under conditions where Hsp90 is inhibited [12, 18]. Consistent with this observation, inhibition of Hsp90 by geldanamycin (GA) followed by a time-dependent immunoprecipitation of overexpressed CRAF^WT^ from *CRAF^-/-^* HEK293T cells revealed a significant increase in Hsp90 binding between 4 h and 8 h of GA exposure, while Hsp70 binding progressively increased from 4 h to 16 h (Fig. S1A). In contrast, under normal overexpression conditions without any treatment, mutant CRAF proteins harboring substitutions at the ‘D’ residue of the DFG motif (D486A, D486G, and D486N) displayed markedly elevated binding to Hsp90, whereas their association with Hsp70 remained largely unchanged (Fig. S1B) [13]. Further analysis showed that both Hsp90α and Hsp90β exhibited significantly increased association with CRAF^WT^ following GA treatment for 4 hours, as well as with CRAF^D486N^ under untreated conditions. These observations indicate that both Hsp90 isoforms exert similarity to regulate CRAF^WT^ during Hsp90 inhibition and on CRAF^D486N^ under basal conditions (Fig. 1A). It has previously been reported that inhibition of Hsp90 destabilizes CRAF^WT^ while simultaneously increasing its association with Hsp90 [12, 13]. To determine whether a similar effect occurs in the case of CRAF^D486N^-which already exhibits strong Hsp90 association-the steady-state level of overexpressed CRAF^D486N^ was monitored in *CRAF^-/-^* cells following time-dependent treatment with GA. At early time points of GA exposure (up to 4-8 h), both WT and D486N proteins showed a similar rate of reduction in steady-state levels. Unexpectedly, with prolonged GA exposure the level of CRAF^D486N^ recovered, whereas CRAF^WT^ continued to decline (Fig. 1Bi & Bii). This divergent behavior suggested distinct chaperone dynamics between the two proteins. Consistent with a stress-responsive effect, Hsp70 binding to D486N gradually increased over time, mirroring the pattern observed for CRAF^WT^ upon GA treatment (Fig. 1C). Notably, D486N exhibited a pronounced surge in Hsp90 binding at an early time point (2-4 h of GA treatment), whereas the increased Hsp90 association with CRAF^WT^ occurred later, between 4 h and 8 h. This little discrimination of early surge in Hsp90 binding to D486N showed a negative correlation with the stability of the mutant protein during GA exposure. Upon prolonged Hsp90 inhibition, Hsp90 binding to D486N stabilized at a plateau or returned to basal levels, which coincided with the recovery of D486N protein levels (Fig. 1C). Importantly, no such changes in D486N stability or chaperone association were observed in DMSO-treated *CRAF^-/-^*cells (Fig. S1C), confirming that these effects were specific to Hsp90 inhibition. To determine whether the transient destabilization of D486N following the surge in Hsp90 binding was accompanied by altered ubiquitination, an in vivo ubiquitination assay was performed. At an early time point of GA exposure (4 h), corresponding to the period immediately following the Hsp90 surge, ubiquitination of D486N was markedly increased. In contrast, during prolonged GA exposure-when Hsp90 dissociated and returned to basal binding levels despite sustained Hsp70 association-the ubiquitination of D486N was reduced (Fig. 1D). Together, these findings suggest that Hsp90 plays a critical role in initiating the degradation of CRAF^D486N^. One possible interpretation is that Hsp90 facilitates the recruitment of an ubiquitin ligase to the mutant CRAF protein or Hsp90 may transmit a signal through the dissociation from client that triggers the degradation process. In either scenario, the transient surge in Hsp90 binding appears to initiate the degradation program and fails to protect this strong client protein once Hsp90 activity is inhibited at early time points.

**Figure 1:**
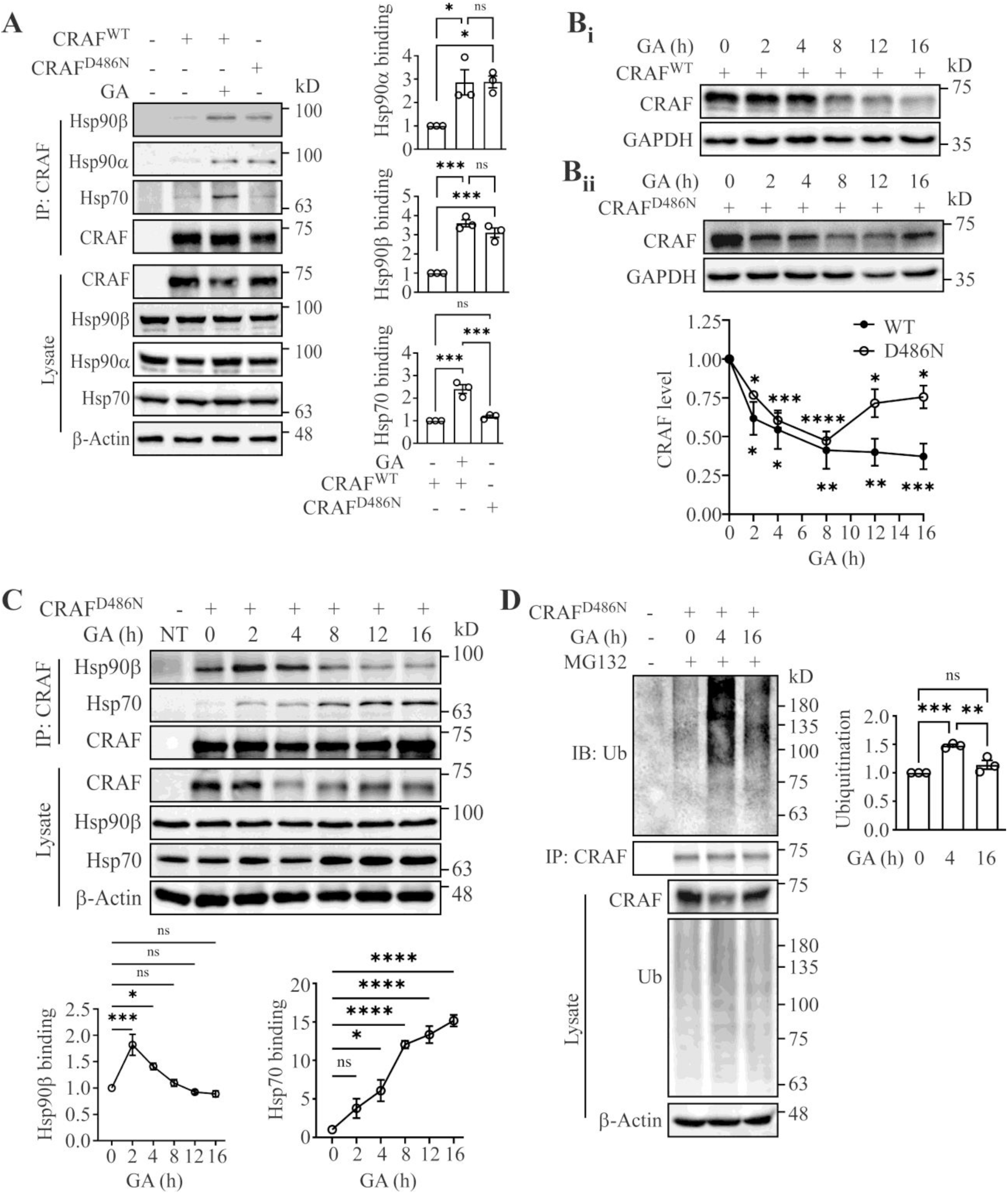
Hsp90 initiates degradation process of the D486N mutant CRAF. **(A)** Interaction of Hsp90α, Hsp90β, and Hsp70 with CRAF^WT^ (∼2 µM geldanamycin for 4h) and CRAF^D486N^ was assessed by immunoprecipitation. CRAF^WT^ and CRAF^D486N^ were overexpressed in *CRAF^-/-^*HEK293T cells for 48h and immunoprecipitated with CRAF antibody. Association of chaperones was observed with their corresponding antibodies. For statistical analysis, one-way ANOVA was performed using GraphPad Prism (n=3). Significance levels: ns (non-significant), \**p* < 0.05, *** *p* < 0.001. **(B)** Expression of overexpressed CRAF^WT^ (Bi) & CRAF^D486N^ (Bii) in *CRAF^-/-^*HEK293T cells was detected by immunoblotting with CRAF antibody following geldanamycin (GA) treatment at the indicated time points (0h to 16h) noted as GA(h). For statistical analysis, one-way ANOVA was performed using GraphPad Prism (n=3). Significance levels: \**p* < 0.05, \*\**p* < 0.01, *** *p* < 0.001, \*\*\*\**p* < 0.0001. **(C)** CRAF^D486N^ was overexpressed in *CRAF^-/-^*HEK293T cells and immunoprecipitated with CRAF antibody at the indicated time points in the presence of geldanamycin. The association of Hsp90β and Hsp70 was evaluated by immunoblotting using their respective antibodies. Statistical analysis, one-way ANOVA, was performed using GraphPad Prism (n=3). Significance levels: ns (non-significant) *p* > 0.05; \**p* < 0.05; \*\*\**p* < 0.001; \*\*\*\**p* < 0.0001. **(D)** CRAF^D486N^ was overexpressed in *CRAF^-/-^* cells, followed by treatment with MG132 (5µM for 4h) and geldanamycin at the indicated time points. CRAF^D486N^ was immunoprecipitated using a CRAF antibody, and endogenous ubiquitination was detected by immunoblotting with a ubiquitin antibody. Statistical analysis was performed using one-way ANOVA in GraphPad Prism (n=3). Significance levels: ns (non-significant) *p* > 0.05; \*\**p* < 0.01; \*\*\**p* < 0.001.

### HECTD3 ubiquitin ligase destabilizes CRAF^D486N^

A prior study by Noble et al., indicated that the ubiquitin ligase CHIP is involved in the degradation of wild-type CRAF (CRAF^WT^), but not in the degradation of the kinase-dead mutant CRAF^D486A^. To further investigate the role of ubiquitin ligases in regulating the selectivity and stability of another similar kinase-dead mutant, D486N, stable shRNA-mediated ubiquitin ligase-silenced *CRAF^-/-^* cell was generated (Fig. S2). To identify the ubiquitin ligase responsible for regulating this kinase-dead mutant, a screening strategy was attempted in which kinase-dead mutants (K375W and S621A, along with D486N) and wild-type (in the absence and presence of GA) were overexpressed in the ubiquitin ligase-silenced cells, and their stability was subsequently examined, were chosen on the basis of their potency to degrade the client of Hsp90 described previously [15, 16, 19–21]. An increase in the level of CRAF^WT^ was observed under CHIP- and Cul5-silenced conditions (Fig. S2Ai and Bi). However, no change in the levels of the mutant proteins was detected under the CHIP-silenced condition (Fig. S2Aii). In contrast, a partial stabilization of the S621A mutant was observed under the Cul5-silenced condition (Fig. S2Bii). No elevation in the levels of wild-type or mutant was detected when UHRF2 or HERC2 were silenced (Fig. S2Ci, Cii, Di, Dii, & Diii). Notably, stabilization of CRAF in cells expressing shHECTD3-1, stabilization of CRAF^WT^ (both in the absence and presence of GA) as well as the kinase-dead mutants D486N and K375W was detected (Fig. 2A, S2Ei & Eii). An additional stable HECTD3-silenced *CRAF^-/-^* cell (shHECTD3-2) was generated, and a similar stabilization phenotype was observed (Fig. 2A) to confirm the previous one. Importantly, no change in the expression level of GFP was detected in the HECTD3-silenced condition, indicating that the observed effect was not due to nonspecific changes in protein expression (Fig. S2Eiii). To determine whether the effect of HECTD3 on D486N occurs through a direct or indirect mechanism, an immunoprecipitation assay was performed. A direct interaction between overexpressed HECTD3 and the D486N mutant of CRAF was observed (Fig. 2B). To further examine whether this interaction influences ubiquitination, the D486N mutant was overexpressed in the presence of HECTD3^WT^ or mutant HECTD3 (HECTD3^DM^). Increased poly-ubiquitination of D486N was detected in the presence of HECTD3^WT^, whereas no change in ubiquitination pattern was observed when HECTD3^DM^ was overexpressed (Fig. 2C). To examine whether recruitment of HECTD3 and D486N changes during time-dependent Hsp90 inhibition, alongside the association of Hsp90 and Hsp70, HECTD3 and D486N were overexpressed in *CRAF^-/-^* and treated with GA in a time-dependent manner. CRAF was immunoprecipitated and the interaction between HECTD3 and D486N gradually increased starting from 2 h of GA treatment and remained elevated even during prolonged GA exposure was observed (Fig. 2D). Notably, this interaction did not decrease despite the reduction in Hsp90 association observed at later time points (after 4 h). This observation suggests that although recruitment of HECTD3 may occur through Hsp70, there is a signal mediated by Hsp90 is likely required for the degradation initiation. If recruitment alone were responsible, a decline in HECTD3 association would have been expected in parallel with the decrease in Hsp90 binding; however, this was not observed (Fig. 2D). An additional observation was that the cellular level of HECTD3 was gradually reduced to a significant extent upon GA treatment (Fig. 2D). This finding suggests that HECTD3 itself may function as an Hsp90 client protein, similar to CRAF^WT^. Such a possibility has also been reported previously [16]. Interestingly, HECTD3 has been reported to be predominantly expressed in heart-derived cell lines (other than many cancer cell lines) and to attenuate hypertrophic responses [22, 23]. Since mutant forms of CRAF are also associated with hypertrophic signaling, the possibility arises that regulation of HECTD3 levels may influence the stability and regulation of mutant CRAF proteins.

**Figure 2:**
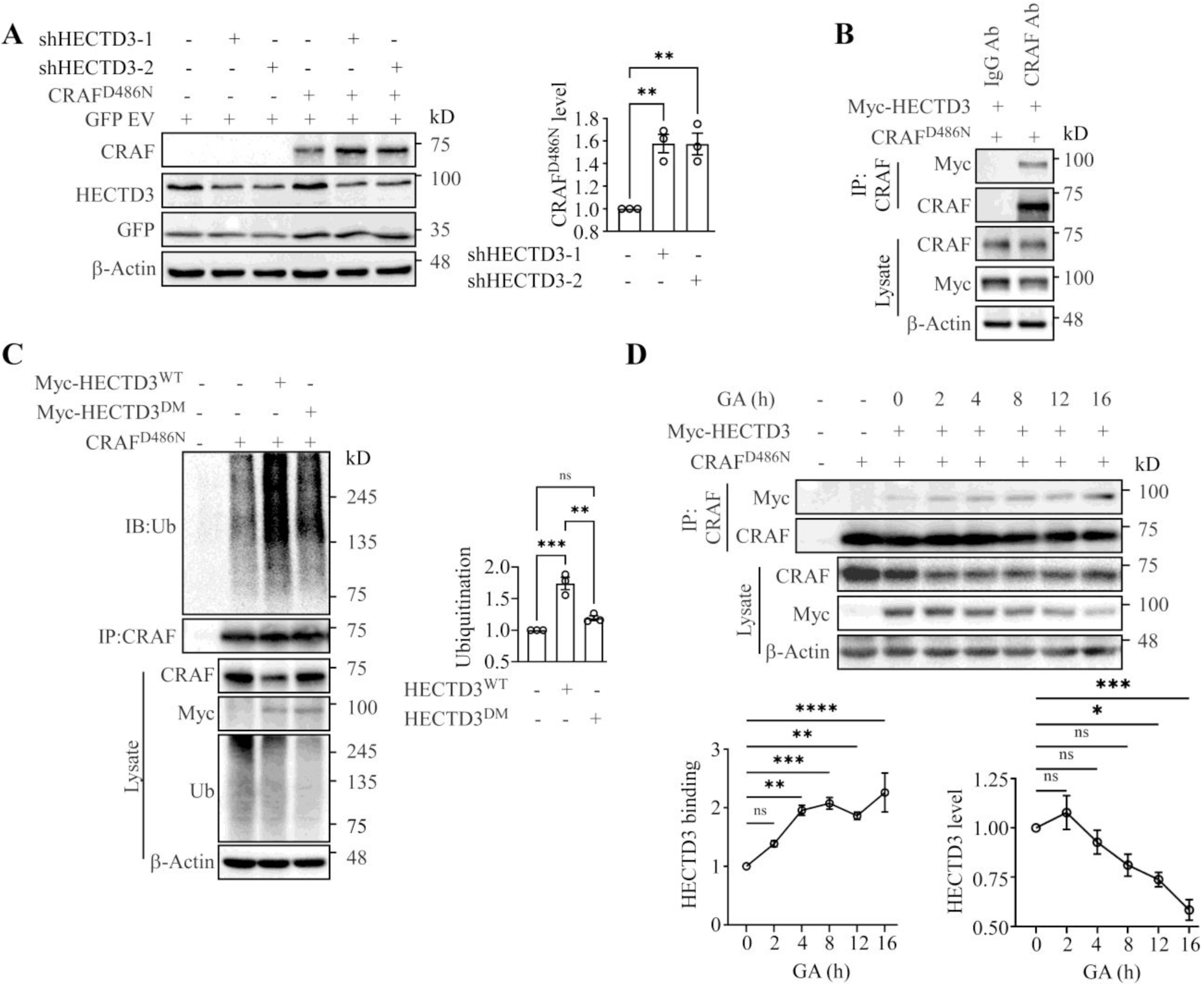
HECTD3-mediated degradation of D486N mutant CRAF. **(A)** CRAF^D486N^ was transfected into shHECTD3-1 and shHECTD3-2 stable *CRAF^-/-^* HEK293T cells. D486N CRAF expression level was detected using a CRAF-specific antibody. Quantification of CRAF expression was performed by normalizing to GFP (empty vector control), shown for CRAF^D486N^. Here, the GFP empty vector served as the transfection control for overexpression plasmids. Quantification was normalized to the GFP/GAPDH ratio and analyzed by Student’s t-test using GraphPad Prism (n=3). Statistical significance: \*\**p* < 0.01. **(B)** Myc-tagged HECTD3 and the CRAF^D486N^ were co-transfected in *CRAF^-/-^*HEK293T cells for 48h. Immunoprecipitation of the CRAF^D486N^ was performed using an anti-CRAF antibody, and the presence of co-immunoprecipitated HECTD3 was confirmed by immunoblotting with an anti-Myc antibody. **(C)** CRAF^D486N^ was overexpressed in *CRAF^-/-^* cells, along with Myc-HECTD3^WT^ or Myc-HECTD3^DM^ followed by treatment with MG132 (5µM for 4h). CRAF^D486N^ was immunoprecipitated using a CRAF antibody, and poly-ubiquitination was detected by immunoblotting with a ubiquitin antibody. Statistical analysis was performed using one-way ANOVA in GraphPad Prism (n=3). Significance levels: ns (non-significant) *p* > 0.05, \*\**p* < 0.01, \*\*\**p* < 0.001. **(D)** CRAF^D486N^ was overexpressed in *CRAF^-/-^*HEK293T cells and immunoprecipitated with CRAF antibody at the indicated time points in the presence of geldanamycin. The association of Myc-HECTD3 was evaluated by immunoblotting using their respective antibodies. Statistical analysis, one-way ANOVA, was performed using GraphPad Prism (n=3). Significance levels: ns (non-significant) *p* > 0.05, \**p* < 0.05, \*\**p* < 0.01, \*\*\**p* < 0.001, \*\*\*\**p* < 0.0001.

### Cul5-mediated destabilization of HECTD3 under stress conditions

During Hsp90 inhibition, Cul5 has been reported to be a major ubiquitin ligase responsible for the degradation of several Hsp90 client proteins, a process that is often facilitated by Hsp70 [16, 19, 24]. Under such conditions, destabilization of HECTD3 was observed (Fig. 2D). Consistent with this observation, the destabilization of both endogenous HECTD3 (Fig. S3Ai) and overexpressed HECTD3 (Fig. S3Aii) was found to be rescued upon silencing of the Cul5 E3 ligase using shCul5. However, no significant change in the basal level of HECTD3 (both endogenous and overexpressed) was detected under this shCul5-mediated silencing condition, likely due to substantial abundance of residual Cul5. Notably, when the level of Cul5 silencing was substantially increased using sgCul5-1 and sgCul5-2, a pronounced rescue of HECTD3 levels was observed (Fig. 3A & B). A similar rescue effect was also observed upon proteasomal inhibition using MG132, indicating that Cul5 functions as one of the regulators of HECTD3 both under GA-treated stress conditions and under normal conditions. Interestingly, sgRNA-mediated Cul5 silencing was associated with a significant reduction in cell size or cellular area compared with WT HEK293T cells, and this phenotype was restored upon overexpression of Myc-tagged Cul5 (Fig. S3B), suggesting a functional role of Cul5 in regulating cellular morphology and cell shape. Since HECTD3 has been reported to be predominantly expressed in heart muscle cells and is known to play a role in attenuation of hypertrophy, the effect of stress conditions on HECTD3 and Cul5 was further examined in cardiomyocyte-derived cell lines [23]. In H9c2 and AC16 cells, destabilization of HECTD3 along with upregulation of Cul5 was observed, in contrast to HEK293T cells, where prominent Cul5 upregulation was not detected upon geldanamycin or other stressogenic treatments, although a decrease in overexpressed HECTD3 levels was still observed (Fig. S3Ci, Cii, & Ciii). Given the prominent upregulation of Cul5 and destabilization of HECTD3 observed in AC16 cells, together with the observed effect of Cul5 on cellular morphology in HEK293T cells, the functional consequence of Cul5 overexpression was examined in AC16 cells through real-time PCR data. It was observed that overexpression of Cul5 led to the upregulation of the mRNA of hypertrophic markers ANP and BNP in the AC16 cell line (Fig. S3D), indicating a potential link between Cul5 activity and hypertrophic signaling [25]. To determine whether the effect of Cul5 on HECTD3 occurs through a direct or indirect mechanism, an immunoprecipitation study was performed in *CRAF^-/-^* cells under basal conditions as well as following treatment with geldanamycin, phenylephrine (PE), and lipopolysaccharide (LPS), since CRAF is a known common interacting partner of both Cul5 and HECTD3. A significant increase in the interaction between Cul5 and HECTD3 was observed upon PE and LPS treatment (Fig. S3E). In addition, increased association of Hsp90, Hsp70, and Cul5 was detected upon GA treatment, suggesting that molecular chaperones may facilitate this interaction process (Fig. 3C). Interestingly, in addition to the designated Cul5 protein band, an additional Cul5 band was detected in the GA-treated samples during the association with HECTD3. This band is likely to represent a post-translationally modified form of Cul5, suggesting the possibility of a well-known regulatory modification of Cul5, most likely its neddylation activation modification [26]. Consistent with this interpretation, the appearance of this additional band was reduced when a neddylation activation modification-deficient Cul5 construct was generated to further investigate the role of Cul5 neddylation in regulating HECTD3. A neddylation-deficient Cul5 mutant (Cul5^K724R^ also Cul5^ΔNRDD8^) was generated, and ubiquitination analysis was performed in sgCul5-1 mediated *Cul5^-/-^*(knock-out) cells (Fig. 3D). As expected, ubiquitination of HECTD3 was increased upon overexpression of Cul5^WT^, whereas no such increase was observed upon overexpression of the Cul5^ΔNRDD8^. These findings indicate that neddylation of Cul5 is required for the regulation of HECTD3 levels (Fig. 3D).

**Figure 3:**
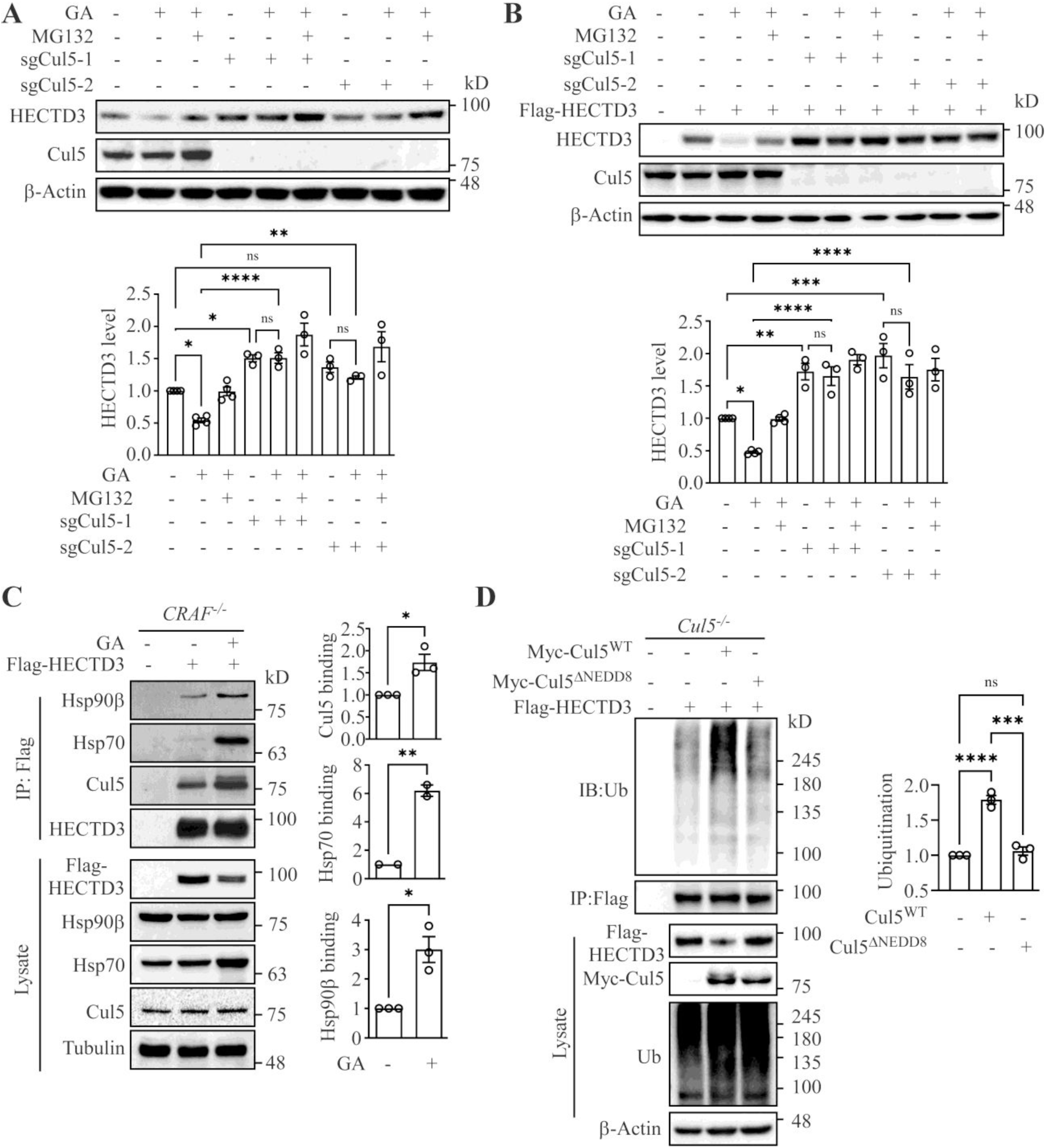
Destabilization of HECTD3 is regulated by Cul5. **(A)** Endogenous HECTD3 level was assessed in Cul5-silenced HEK293T polyclonal (sgCul5-1 and sgCul5-2) cells in absence and presence of GA (2µM) and MG132 (5µM), as indicated for 16h in HEK293T cell. HECTD3 and Cul5 expression was assessed via western blot using an anti-HECTD3 and anti-Cul5 antibody respectively. Statistical analysis was performed using one-way ANOVA in GraphPad Prism (n=3). Significance levels: ns (non-significant) *p* > 0.05, \**p* < 0.05, \*\**p* < 0.01, \*\*\*\**p* < 0.0001. **(B)** Flag-tagged HECTD3 was overexpressed in Cul5-silenced HEK293T polyclonal (sgCul5-1 and sgCul5-2) cells for 48h in absence and presence of GA (µM) and MG132 (5µM), for 16h as indicated. HECTD3 and Cul5 expression was assessed via western blot using an anti-Flag and anti-Cul5 antibody respectively. Statistical analysis was performed using one-way ANOVA in GraphPad Prism (n=3). Significance levels: ns (non-significant) *p* > 0.05, \**p* < 0.05, \*\**p* < 0.01, \*\*\**p* < 0.001, \*\*\*\**p* < 0.0001. **(C)** Flag-tagged HECTD3 was overexpressed in *CRAF^-/-^* HEK293T cells in the absence and presence of GA treatment for 16h. HECTD3 was immunoprecipitated using an anti-Flag antibody, and the co-immunoprecipitated proteins Cul5, Hsp70, and Hsp90β were detected using their respective antibodies and measured by immunoprecipitation band aspect ratio. Statistical significance was determined using a Student’s t-test, GraphPad Prism. Notation: \**p* < 0.05, \*\**p* < 0.01. **(D)** Flag-tagged HECTD3 was overexpressed alone or co-expressed with either wild-type Myc-Cul5 (Cul5^WT^) or a neddylation-deficient mutant (Myc-Cul5^K724R^/Myc-Cul5^ΔNEDD8^) in Cul5 knockout (sgCul5-1) polyclonal HEK293T cells followed by treatment with MG132 (5µM for 4h). HECTD3 was immunoprecipitated using Flag antibody and poly-ubiquitination of overexpressed HECTD3 was assessed. Statistical significance was determined using GraphPad Prism (n=3). Significance level: ns (non-significant) *p* > 0.05, \*\*\**p* < 0.001, \*\*\*\**p* < 0.0001.

### Neddylation activation of Cul5 stabilizes CRAF^D486N^

As stress conditions were found to induce destabilization of HECTD3 and upregulation of Cul5 levels (Fig. 3), the status of CRAF^D486N^-identified as a substrate of the HECTD3 ubiquitin ligase-was examined under this conditions. For this purpose, *CRAF-Cul5* double knockout HEK293T cells (*CRAF^-/-^Cul5^-/-^*) were generated (Fig. S4A) to eliminate the traces of endogenous Cul5 inside the cell. Wild-type Cul5 (Cul5^WT^) and neddylation activation modification-deficient Cul5 (Cul5^ΔNEDD8^) were then overexpressed in a dose-dependent manner (to imitate the upregulated Cu5 conditions) together with CRAF^WT^ or CRAF^D486N^ in these *CRAF^-/-^Cul5^-/-^* HEK293T cells, and the levels of CRAF (WT as well as D486N) were subsequently assessed. Upon Cul5^WT^ overexpression, a gradual decrease in the level of CRAF^WT^ was observed (Fig. S4Bi). In contrast, no significant changes in the level of CRAF^WT^ were detected upon overexpression of Cul5^ΔNEDD8^ (Fig. S4Bii). Interestingly, a distinct effect was observed for the CRAF^D486N^. A significant gradual increase in the level of CRAF^D486N^ was detected upon dose-dependent overexpression of Cul5^WT^, whereas no change in the level of CRAF^D486N^ was observed in the presence of Cul5^ΔNEDD8^ overexpression (Fig. 4Ai & Aii). To investigate the mechanism by which Cul5 stabilizes CRAF^D486N^, the association between overexpressed HECTD3 and the D486N mutant was examined in the presence of Cul5^WT^ or Cul5^ΔNEDD8^. For this experiment, HECTD3 and CRAF^D486N^ were co-overexpressed in *CRAF^-/-^Cul5^-/-^* cells, followed by immunoprecipitation of D486N (Fig. 4C). It was observed that the association between HECTD3 and CRAF^D486N^ was decreased in the presence of both Cul5^WT^ and Cul5^ΔNEDD8^ (Fig. 4B). Consistent with this observation, a decreased poly-ubiquitination level of CRAF^D486N^ was detected in the presence of Cul5^WT^, whereas no such change was observed in the presence of the Cul5^ΔNEDD8^ (Fig. 4C). These findings indicate that Cul5 (WT or neddylation deficient mutant) contributes to the dissociation of HECTD3 from CRAF^D486N^, and that the neddylation activation modification of Cul5 further reduces the ubiquitination level of CRAF^D486N^. Since CRAF^D486N^ is a hypertrophic mutant associated with RASopathies, the physiological consequences of its stabilization were subsequently examined. To this end, the D486N mutant of CRAF was overexpressed in the AC16 cell line, and hypertrophic signaling was assessed by examining the mRNA expression of the hypertrophic marker genes ANP and BNP by real-time PCR. However, activation of these hypertrophic signaling genes was not observed in AC16 cells upon overexpression of CRAF^D486N^ (Fig. S4C), a finding that is consistent with previous observations [27, 28]. Consequently, further investigation was initiated to examine the interplay between Cul5 and HECTD3 and its physiological consequences in other hypertrophic mutants of CRAF (S257L and L613V) in both HEK293T and AC16 cells.

**Figure 4:**
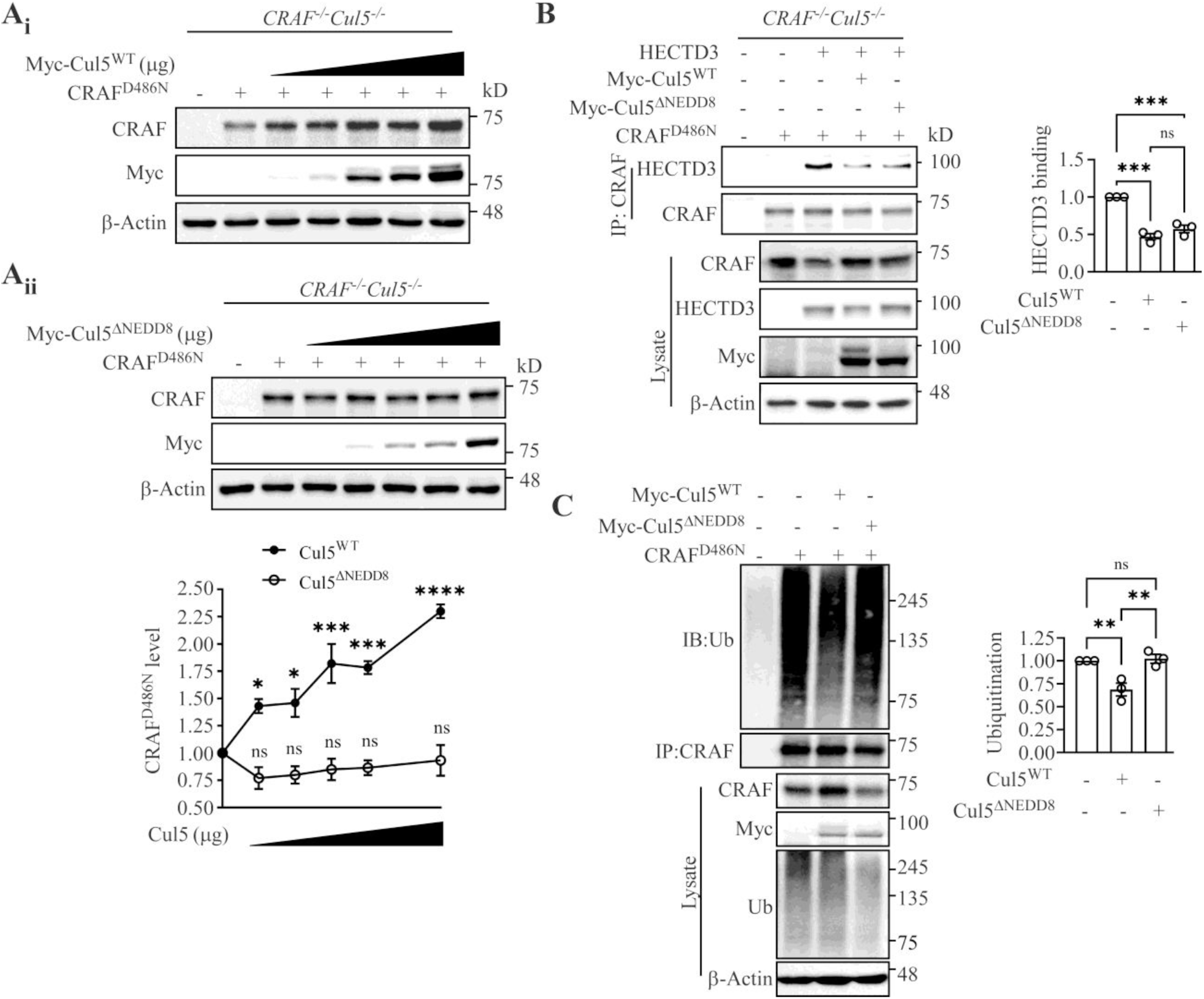
Cul5 overexpression stabilizes D486N mutant CRAF. **(A)** CRAF^D486N^ was overexpressed in *CRAF^-/-^Cul5^-/-^* double knockout (DKO) HEK293T cells, Cul5^WT^ and Cul5^ΔNEDD8^ were also overexpressed along with CRAF^D486N^ in a dose dependent overexpression manner (in µg) at a ratio from 0.2 to 1 (CRAF/Cul5). The level of CRAF was evaluated by immunoblotting using CRAF antibody. Statistical analysis, one-way ANOVA, was performed using GraphPad Prism (n=3). Significance levels: ns (non-significant) *p* > 0.05; \**p* < 0.05; \*\*\**p* < 0.001; \*\*\*\**p* < 0.0001. **(B)** CRAF^D486N^, Flag-HECTD3, Myc-Cul5^WT^ and Myc-Cul5^ΔNEDD8^ were overexpressed in *CRAF^-/-^Cul5^-/-^*double knockout (DKO) cells for 48h as indicated in the figure. Immunoprecipitation of the CRAF^D486N^ was performed using an anti-CRAF antibody, and the presence of co-immunoprecipitated HECTD3 was confirmed by immunoblotting with an anti-HECTD3 antibody. Statistical analysis, one-way ANOVA, was performed using GraphPad Prism (n=3). Significance levels: ns (non-significant) *p* > 0.05, \*\*\**p* < 0.001. **(C)** CRAF^D486N^ was overexpressed alone or co-expressed with either wild-type Myc-Cul5 (Cul5^WT^) or a neddylation-deficient mutant (Myc-Cul5^K724R^/Myc-Cul5^ΔNEDD8^) followed by treatment with MG132 (5µM for 4h) in *CRAF^-/-^Cul5^-/-^* double knockout (DKO) cells. CRAF was immunoprecipitated using CRAF antibody and poly-ubiquitination of overexpressed CRAF was assessed. Statistical significance was determined using GraphPad Prism (n=3). Significance level: ns (non-significant) *p* > 0.05; \*\**p* < 0.01.

### Hypertrophic mutant CRAF shares partially similar regulatory behavior

Following the negative result obtained with the D486N mutant in hypertrophic signaling assays, additional hypertrophic CRAF mutants, S257L and L613V, were examined similarly. These mutants were overexpressed in *CRAF^-/-^Cul5^-/-^* cells, followed by gradual overexpression of Cul5^WT^ or Cul5^ΔNEDD8^ constructs. Under these conditions, no change in the level of the CRAF^L613V^ was observed in the presence of either Cul5^WT^ or Cul5^ΔNEDD8^ (Fig. S5Ai & Aii). In contrast, a significant increase in the level of the CRAF^S257L^ was detected upon gradual overexpression of Cul5^WT^, whereas no alteration was observed in the presence of Cul5^ΔNEDD8^ (Fig. 5Ai & Aii). As, CRAF^S257L^ showed similar behavior with CRAF^D486N^, the ubiquitination status of this mutant was subsequently examined. It was observed that the poly-ubiquitination level of CRAF^S257L^ was significantly decreased in the presence of Cul5^WT^, while no change in ubiquitination was detected upon overexpression of the Cul5^ΔNEDD8^ (Fig. S5B). To find the reason behind the different behavior between CRAF^S257L^ and CRAF^L613V^ upon Cul5 overexpression, whether these hypertrophic mutants are regulated in a manner similar to CRAF^D486N^ and whether they represent substrates of HECTD3, the levels of CRAF^S257L^ and CRAF^L613V^ were examined in the absence and presence of HECTD3 silencing using shHECTD3-1 and shHECTD3-2. Stabilization of CRAF^S257L^ was observed upon HECTD3 silencing, whereas no significant alteration in the level of CRAF^L613V^ was detected (Fig. 5B); and as a followed experiment the regulation of CRAF^S257L^ by HECTD3 through ubiquitination was checked with WT or mutant HECTD3. As expected, an increase in poly-ubiquitination of CRAF^S257L^ was observed in the presence of HECTD3^WT^, whereas no change in ubiquitination was detected when HECTD3^DM^ was overexpressed (Fig. S5C). The discriminating observation from early experiment (Fig. 5B) triggered to determine the association of HECTD3 with CRAF^L613V^ probably the key to regulate the mutants. Surprisingly, like CRAF^S257L^, CRAF^L613V^ can also bind with HECTD3 (Fig. 5C). To understand why CRAF^L613V^ exhibits regulatory behavior distinct from that of CRAF^S257L^, the association of these mutants with molecular chaperones was observed taking CRAF^WT^ as a control. It was observed that CRAF^S257L^ displayed increased association with both Hsp90β and Hsp70. In contrast, CRAF^L613V^ exhibited increased association with Hsp70 but reduced association with Hsp90β (Fig. 5D). To further examine the role of chaperones in HECTD3 recruitment, Hsp90β and Hsp70 were individually overexpressed together with CRAF^L613V^ and HECTD3. Under these conditions, a significant increase in the association between CRAF^L613V^ and HECTD3 was observed in both cases; however, the interaction was more pronounced under Hsp70 overexpression conditions (Fig. S5D). These observations suggest that both Hsp90β and Hsp70 are capable of recruiting HECTD3. This may explain why CRAF^L613V^ can interact with HECTD3 but is not effectively regulated by Hsp90β due to its relatively weak association with this chaperone. To further investigate the specific contribution of Hsp90, a CRAF-Hsp90β double knockout (*CRAF^-/-^Hsp90β^-/-^*) HEK293T cell line was generated. In this system, the association of HECTD3 substrate CRAF^S257L^ with Hsp90α was examined in the absence and presence of Hsp90β (both endogenous and overexpressed). CRAF^S257L^ was overexpressed in the *CRAF^-/-^Hsp90β^-/-^* and subsequently immunoprecipitated. An increased association of Hsp90α was observed in the absence of Hsp90β, and when Hsp90β expression was restored through overexpression, no significant change in Hsp90α association was detected (Fig. S5E). These observations indicate that further overexpression of Hsp90β does not significantly alter the association of Hsp90α, which is already associated with CRAF^S257L^. Under these conditions, the association of HECTD3 was further examined in the absence and presence of Hsp90β (wild-type Hsp90β, Hsp90β^WT^, as well as ATPase mutant Hsp90β, Hsp90β^DN^). It was observed that HECTD3 association with CRAF^S257L^ was significantly increased in the presence of Hsp90β^WT^, where stronger Hsp90β association was also detected. In contrast, the Hsp90β^DN^ showed reduced association with CRAF^S257L^, and lower recruitment of HECTD3 to CRAF^S257L^ (Fig. 5E). Collectively, these observations indicate that Hsp70 can recruit HECTD3 to mutant CRAF proteins; however, the ATPase activity of Hsp90 appears to provide an additional signal required for efficient recruitment and degradation. Even in the absence of Hsp90, recruitment by Hsp70 can occur, but without the ATPase activity of Hsp90, the degradation by HECTD3 association isn’t triggered, suggesting that Hsp90-mediated signaling contributes to degradation initiation process. To assess whether overexpression of CRAF^S257L^ induces cellular stress prior to evaluating its physiological consequences, the association between Cul5 and HECTD3 was examined in the absence and presence of the CRAF^S257L^. An increased association between Cul5 and HECTD3 was observed in the presence of CRAF^S257L^ (Fig. S5F), indicating that stress conditions induced by hypertrophic mutant CRAF can promote enhanced binding of Cul5 to HECTD3.

**Figure 5:**
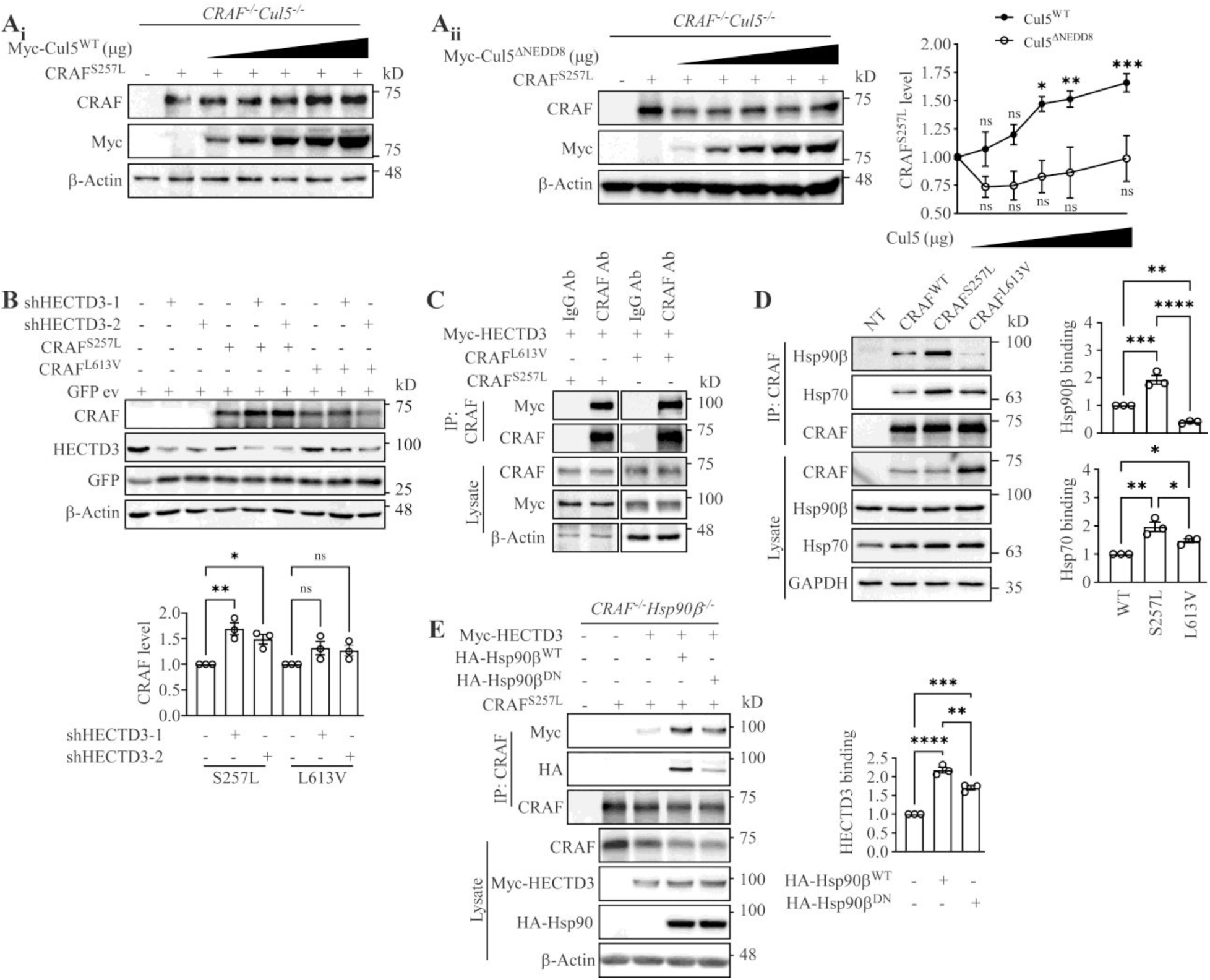
Regulation of hypertrophic mutant S257L by Hsp90 and HECTD3. **(A)** CRAF^S257L^ was overexpressed in *CRAF^-/-^Cul5^-/-^* cells, Cul5^WT^ and Cul5^ΔNEDD8^ were also overexpressed along with CRAF^S257L^ in a dose dependent overexpression manner (in µg) at a ratio from 0.2 to 1 (CRAF/Cul5). The level of CRAF was evaluated by immunoblotting using CRAF antibody. Statistical analysis, one-way ANOVA, was performed using GraphPad Prism (n=3). Significance level: ns (non-significant) *p* > 0.05; \**p* < 0.05, \*\**p* < 0.01, \*\*\**p* < 0.001. **(B)** CRAF^S257L^ and CRAF^L613V^ were transfected into shHECTD3-1 and shHECTD3-2 stable *CRAF^-/-^* HEK293T cells. Mutant CRAF expression level was detected using a CRAF-specific antibody. Quantification of CRAF expression was performed by normalizing to GFP (empty vector control), shown for mutant CRAFs. Here, the GFP empty vector served as the transfection control for overexpression plasmids. Quantification was normalized to the GFP/GAPDH ratio and analyzed by Student’s t-test using GraphPad Prism (n=3). Significance level: ns (non-significant) *p* > 0.05; \**p* < 0.05, \*\**p* < 0.01. **(C)** Myc-tagged HECTD3 was co-transfected with CRAF^S257L^ or CRAF^L613V^ in *CRAF^-/-^*HEK293T cells for 48h. Immunoprecipitation of the mutant CRAF was performed using an anti-CRAF antibody, and the presence of co-immunoprecipitated HECTD3 was confirmed by immunoblotting with an anti-Myc antibody. **(D)** CRAF^WT^, CRAF^S257L^, and CRAF^L613V^ were overexpressed in *CRAF^-/-^*HEK293T cells and immunoprecipitated with CRAF antibody. The association of Hsp90β and Hsp70 was evaluated by immunoblotting using their respective antibodies. Statistical analysis, one-way ANOVA, was performed using GraphPad Prism (n=3). Significance level: ns (non-significant) *p* > 0.05; \**p* < 0.05, \*\**p* < 0.01, \*\*\**p* < 0.001, \*\*\*\**p* < 0.0001. **(E)** Association of CRAF^S257L^, HECTD3 and Hsp90β was assessed by co-immuniprecipitation. CRAF^S257L^, HA-Hsp90β^WT^, HA-Hsp90β^DN^ and Myc-HECTD3 were overexpressed/co-overexpressed in *CRAF^-/-^Hsp90β^-/-^* double knockout (DKO) HEK293T cells for 48h as indicated in the figure. Immunoprecipitation of the CRAF^S257L^ was performed using an anti-CRAF antibody, and the presence of co-immunoprecipitated HECTD3 & Hsp90β was confirmed by immunoblotting with an anti-Myc & anti-HA antibody respectively. Statistical analysis, one-way ANOVA, was performed using GraphPad Prism (n=3). Significance levels: \*\**p* < 0.01, \*\*\*\**p* < 0.0001.

### CRAF^S257L^ induced hypertrophy is synergistically enhanced by neddylation-activated Cul5 overexpression

To examine the hypertrophic effect of the S257L and to determine how it is regulated by HECTD3 and Cul5 in cardiomyocytes derived cell, CRAF^S257L^ was overexpressed in the AC16 cell line together with HECTD3^WT^, HECTD3^DM^, Cul5^WT^, or Cul5^ΔNEDD8^. The protein level of S257L was then assessed by western blot analysis. A significant decrease in the level of CRAF^S257L^ was observed upon overexpression of HECTD3^WT^, whereas no significant change in CRAF^S257L^ levels was detected in the presence of HECTD3^DM^ (Fig. 6A). In contrast, upon overexpression of Cul5^WT^, an increase in the level of CRAF^S257L^ was observed, as expected, while no change in CRAF^S257L^ levels was detected upon overexpression of the Cul5^ΔNEDD8^ (Fig. 6A). Consistent with these observations, expression of the hypertrophic marker genes ANP and BNP was found to be upregulated under the conditions of CRAF^S257L^ overexpression, as anticipated. However, this hypertrophic effect was markedly reduced when HECTD3 was overexpressed (Fig. 6B & C). Furthermore, as Cul5 had previously been shown to exhibit a hypertrophic effect (Fig. 3), the expression of hypertrophic genes was examined in the context of combined CRAF^S257L^ and Cul5^WT^ overexpression. Under these conditions, ANP and BNP mRNA expression were found to be upregulated in a synergistic manner when CRAF^S257L^ and Cul5^WT^ were co-overexpressed. In contrast, no significant change in hypertrophic gene expression was observed when CRAF^S257L^ was co-expressed with Cul5^ΔNEDD8^ (Fig. 6B & C). Collectively, these findings demonstrate that hypertrophic mutant CRAF proteins are regulated by the ubiquitin ligases HECTD3 and Cul5, and that modulation of these regulatory pathways correspondingly influences the hypertrophic response.

**Figure 6:**
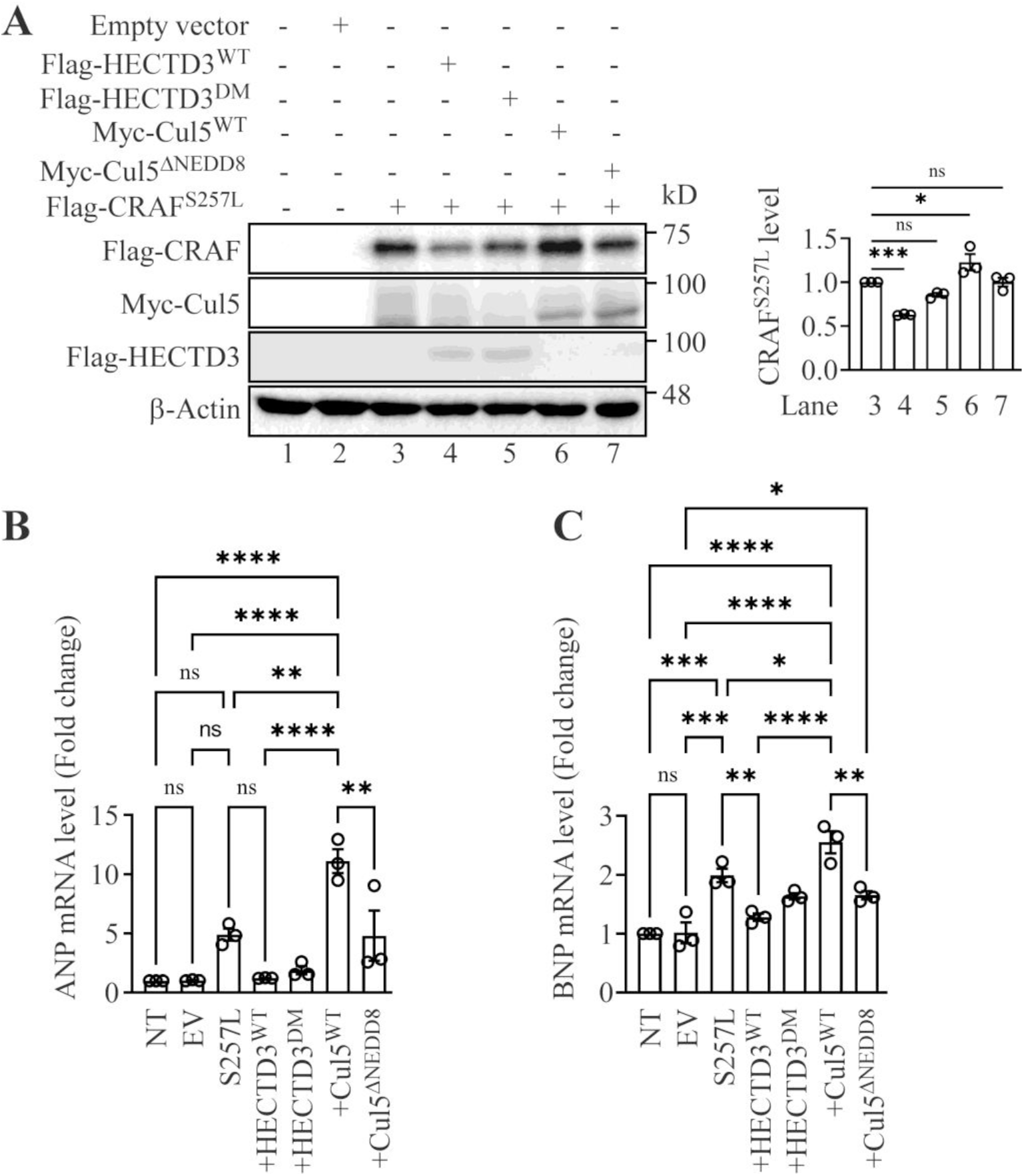
Synergistic effect of S257L and Cul5 in hypertrophy. **(A)** Flag-tagged CRAF^S257L^ was overexpressed alone or co-expressed with wild-type Flag-HECTD3 (HECTD3^WT^) or a catalytic inactive mutant (HECTD3^DM^) or wild-type Myc-Cul5 (Cul5^WT^) or a neddylation-deficient mutant (Myc-Cul5^ΔNEDD8^) in AC16 cells. Expression of S257L, Cul5, HECTD3 and the loading control β-actin was evaluated using anti-Flag, anti-Myc, anti-Flag and anti-β-actin antibodies, respectively. Statistical significance was determined using GraphPad Prism. Significance level: ns (non-significant) *p* > 0.05; \**p* < 0.05, \*\*\**p* < 0.001. **(B)** Similarly like (A), AC16 cells were transfected and the mRNA was isolated and samples were subjected to real time PCR with ANP primers taking Beta actin as control gene. Statistical significance was determined using GraphPad Prism. Significance level: ns (non-significant) *p* > 0.05; \*\**p* < 0.01, \*\*\*\**p* < 0.0001. **(C)** Similarly like (B), AC16 cells were transfected and the mRNA was isolated and samples were subjected to real time PCR with BNP primers taking Beta actin as control gene. Statistical significance was determined using GraphPad Prism. Significance level: ns (non-significant) *p* > 0.05; \**p* < 0.05, \*\**p* < 0.01, \*\*\**p* < 0.001, \*\*\*\**p* < 0.0001.

**Figure 7:**
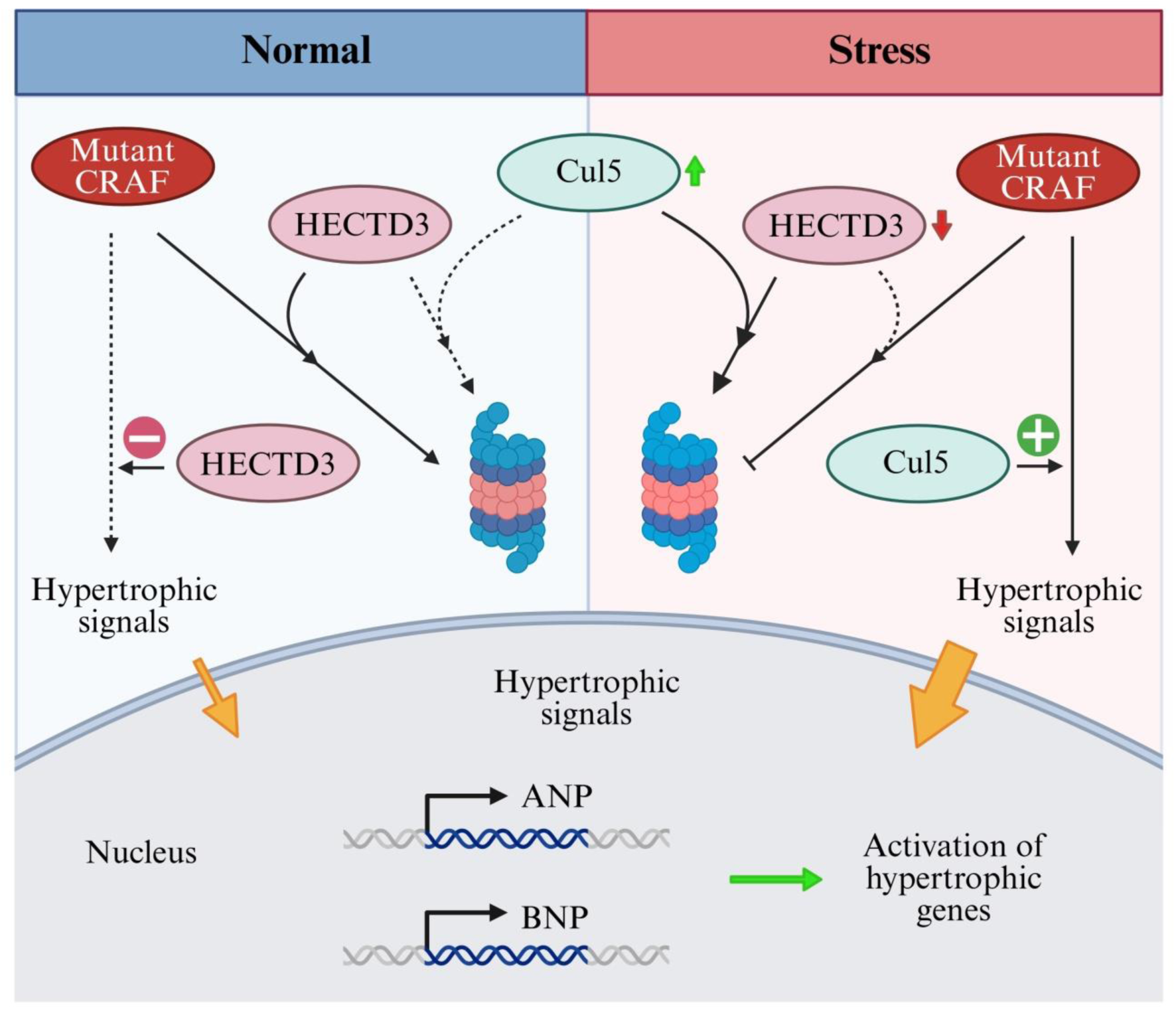
Schematic diagram of mutant CRAF regulation by Cul5-HECTD3 axis. Under physiological conditions, mutant CRAF is degraded via HECTD3-mediated proteasomal pathways and HECTD3 is subjected to minimal degradation by Cul5 in this condition. Hypertrophic signaling to the nucleus is attenuated due to efficient regulation of mutant CRAF by HECTD3. Under stress conditions, HECTD3 is increasingly degraded by Cul5, leading to reduced HECTD3 abundance; mutant CRAF degradation is diminished, and stress-induced upregulation of Cul5 promotes hypertrophic signalling, synergistically with mutant CRAF-mediated hypertrophy, resulting in enhanced transmission of hypertrophic signals to the nucleus.

## Discussion

The regulation of CRAF kinase by ubiquitin ligases is crucial for maintaining proteostasis, particularly in the context of disease-associated mutants. In this study the systematic investigation of the roles of ubiquitin ligases in the degradation of mutant CRAF, providing insights into the distinct mechanisms governing their stability. Our findings reveal a multilayered regulatory axis involving Hsp90-dependent degradation initiation, HECTD3-mediated ubiquitination, and Cul5-dependent control of HECTD3 stability, which together determine the fate of specific CRAF mutants.

CRAF mutants drive pathological signaling in cancers and developmental disorders including Noonan and LEOPARD syndrome [29], understanding their degradation mechanisms is crucial. The regulation of CRAF stability is a highly orchestrated process involving the interplay of molecular chaperones and ubiquitin ligases. Our study dissects this multi-layered regulation in the CRAF mutants, with particular emphasis on chaperone involvement and the mutation-specific nuances that dictate CRAF’s fate within the cell. This highlights ubiquitin ligase, HECTD3 or Cul5 as a potential therapeutic target for diseases linked to CRAF^D486N^ and CRAF^S257L^, where their upregulation or mitigation of activation could regulate mutant CRAF levels. The stabilization of CRAF^D486N^ under prolonged Hsp90 inhibition (GA treatment) further supports the role of chaperone-dependent ubiquitination initiation in CRAF regulation. Since Hsp90 inhibition disrupts wild-type CRAF folding, leading to the degradation by HECTD3[30][29][29][28], our findings suggest that HECTD3 may function similarly in degrading metastable CRAF mutants under physiological condition and also cooperates with Hsp70/Hsp90 in to recognize its substrate.

Initial work by Demand et al. (2001) demonstrated that wild-type (WT) CRAF undergoes degradation facilitated by Hsp70 and its co-chaperone Bag1, positioning Hsp70 as a central player in CRAF proteostasis [15]. However, Noble et al. (2008) later reported that kinase-dead mutants such as D486A bypass this Hsp70-Bag1 pathway, indicating that mutations in CRAF alter its interaction landscape and degradation route [13]. Building on these findings, our immunoprecipitation experiments (Fig. 1) revealed that while CRAF^WT^ shows increased binding to Hsp90 and Hsp70 upon geldanamycin (GA) treatment, kinase-dead mutants, including D486A, D486G, and D486N, exhibit constitutively high affinity for Hsp90 but unaltered binding to Hsp70. This suggests that these mutants adopt metastable conformations that render them more dependent on Hsp90 in basal conditions, because of their impaired kinase folding or ATP binding, which disrupts their native stability [31].

To link chaperone association with functional degradation signals, we assessed ubiquitination of CRAF^D486N^ across early and late time point’s post-GA treatment (Fig. 1). The results showed ubiquitination peaked during early destabilization (4h) and returned to baseline during late stabilization (12h). Interestingly, this ubiquitination mediated destabilization occurred only during the early phase of Hsp90 inhibition, after which the mutant protein gradually recovered despite continued Hsp70 association. This pattern supports a model in which Hsp90 is essential not only for chaperoning CRAF but also for actively governing its ubiquitination, likely by stabilizing a ubiquitination-competent conformation through directly recruiting ubiquitin ligases or by leaving the client, failing to refold the metastable form to stable folded conformation. The transient increase in Hsp90 binding may represent a degradation-priming step that facilitates recruitment of degradation machinery to the client protein. Reduced ubiquitination but unaltered ubiquitin ligase engagement at later time points suggests that either the conformational changes, or the chaperone switching to Hsp70 stabilized the client. These temporal dynamics of chaperone influence on CRAF stability, is supported by Hsp90 inhibition on CRAF^D486N^. ATPase inhibition of Hsp90 stabilized the mutant, whereas high levels of GA treatment destabilized it initially, suggests that CRAF^D486N^, which already has high Hsp90 affinity, and may be sensitive to changes in chaperone equilibrium. Additionally, CRAF^L613V^ can interact with HECTD3, yet its levels remain unchanged upon HECTD3 silencing. This paradox can be resolved by considering its poor binding to Hsp90, suggesting that efficient ubiquitin ligase function may require not just interaction but also chaperone-mediated signal for ubiquitination. Overexpression of Hsp90 and Hsp70 confirmed that both chaperones are capable of recruiting HECTD3, supports dual role of Hsp90: (1) it serves as a recruiter and (2) it provides a conformational signal or ‘permission to degrade’ event that permits the initiation of degradation by probably checking the health of client and leaving client’s in its unchanged form after failing to refold. This aligns with existing literature describing chaperones not just as carriers of unfolded proteins but as active determinants that determine substrate fate based on folding status, ATPase cycling, and co-chaperone interactions. Therefore, HECTD3 recruitment is insufficient to start degradation as it was observed that S257L having low binding with dominant negative form of Hsp90 (mutant ATPase domain) when the background degradative pressure by Hsp90α and Hsp70 is constant and the ATPase domain of Hsp90β is only degradation initiation determining factor observed in *CRAF^-/-^Hsp90β^-/-^*; chaperone-mediated conformational change likely represents a crucial checkpoint.

Such a biphasic response highlights the importance of temporal resolution when interpreting the effects of chaperone modulators. It suggests that therapeutic interventions aimed at modulating chaperone function may have complex, time-sensitive outcomes depending on the mutational status and binding dynamics of the target protein.

The Cullin-RING ligase Cul5 has emerged as a critical regulator of protein quality control, especially under cellular stress conditions that impair protein folding and chaperone function. Cul5 is functionally coupled with molecular chaperones Hsp90 and Hsp70, and is activated via neddylation, a post-translational modification essential for its E3 ligase activity [24]. How Cul5, in its active form, influences the stability of HECTD3 and consequently modulates the degradation of the hypertrophic CRAF was a question of research because HECTD3 attenuates hypertrophy and can destabilized upon Hsp90 inhibition. This vulnerability of HECTD3 suggests it functions like an Hsp90 client-like regulator, whose stability is controlled by the Cul5 and chaperone system. It was also observed stressogenic factors altered the Cul5 expression, possibly the reason which is responsible for HECTD3 degradation, was confirmed by Cul5 silencing rescued endogenous and exogenous HECTD3 from degradation in GA-treated cells, also decreasing the size of the cell, strongly implicating Cul5 in the turnover of HECTD3 under such conditions. To determine whether this regulation involves CRAF dependent effect, we expressed Flag-tagged HECTD3 in *CRAF^-/-^* cells to eliminate CRAF as a potential mediator under Hsp90-inhibited conditions. Co-immunoprecipitation showed enhanced binding of HECTD3 to Cul5, Hsp90, and Hsp70 after GA treatment, reinforcing the hypothesis that both chaperones facilitate substrate recognition and transfer to Cul5 during stress-induced degradation. Cul5, like other Cullin family members, requires neddylation for its activation and substrate ubiquitination. To assess whether this post-translational modification governs HECTD3 degradation, we compared the effects of Cul5^WT^, reduced HECTD3 protein levels, whereas Cul5^ΔNEDD8^ had no such effect. This clearly establishes that neddylation is essential for Cul5 to functionally target HECTD3 for degradation.

Having established that Cul5 mediates the stress-induced degradation of HECTD3, we sought to determine how this regulatory axis influences the stability of mutant CRAF, particularly the hypertrophic mutants observed to be degraded via HECTD3. Our hypothesis was that Cul5 overexpression would relieve degradative pressure on mutant CRAF which is being degraded through HECTD3, resulting in the mutant’s stabilization, because overexpression of Cul5 triggers the hypertrophic effect in AC16 cells.

This designing of a double knockout (DKO) cell line (*CRAF^-/-^Cul5^-/-^*), allowing precise reconstitution of Cul5 and mutant CRAF and eliminates background interactions and provides a clean system for assessing Cul5-dependent effects on CRAF expression. Upon reintroducing Cul5^WT^ or Cul5^ΔNEDD8^ alongside either CRAF^WT^, or CRAF^D486N^, and CRAF^S257L^ we observed striking differences. While Cul5^WT^ reduced the expression of CRAF^WT^, likely through its known role in degrading Hsp90 clients, it stabilized CRAF^D486N^ and CRAF^S257L^ an effect not seen with Cul5^ΔNEDD8^. This dichotomy suggests that Cul5’s effect on CRAF is indirect and mutation-specific, depending on whether HECTD3 is also present and functional. We interpret this as evidence of a competitive degradation mechanism: in the presence of active Cul5, HECTD3 is degraded, lifting the suppression on D486N, which would otherwise be targeted by HECTD3. In contrast, WT CRAF, being a canonical Hsp90 client, is targeted directly by Cul5. This observation is crucial, as it reveals that the fate of mutant proteins such as CRAF^D486N^ and S257L can be modulated by manipulating the stability of their upstream ubiquitin ligase regulators.

These results together suggest a model wherein chaperone acts as a master regulator of both clients and the chaperone-associated ubiquitin ligases. Under stress conditions, such as Hsp90 inhibition, Cul5 is recruited to client-chaperone complexes. This model aligns with broader proteostasis principles, where stress-responsive ubiquitin ligases function not just to degrade misfolded proteins but also to reshape the cellular degradation landscape by removing or modulating other ligases. Such multi-layered regulation ensures that the cell can dynamically adjust protein quality control mechanisms in response to fluctuations in folding capacity, stress, or mutations of proteins.

## Abbreviations

CHIP: C-terminus of Hsc70-interacting protein
CUL: Cullin
HECT: Homologous to the E6-AP carboxyl terminus
HSP: Heat shock protein
HERC: HECT-domain and the RCC1-like domain containing protein
MAPK: Mitogen-activated protein kinase
NEDD8: Neural precursor cell expressed developmentally downregulated protein 8
RAF: Rapidly accelerated fibrosarcoma
UHRF2: Ubiquitin-like with PHD and RING finger domain 2

## Author contribution

SR and AKM conceived the idea and designed the experiments; SR performed most of the experiments; SM contributed in some experiments; SR and AKM analyzed the results; SR wrote the manuscript. All the authors reviewed and corrected the manuscript.

## Data Availability Statement (DAS)

The data represented in the manuscript are available within the article and its supplementary materials which can be shared upon request.

## Acknowledgement

The authors would like to sincerely thank Dhandapany Perundurai for generously providing the CRAF plasmids. We are also grateful to Ashraf Yusuf Rangrez and Norbert Frey for sharing the HECTD3 plasmid, and to Biswanath Maity for kindly providing the AC16 cell line. We extend our sincere appreciation to Barun Mahata, Shibjyoti Debnath, and Abhishek Sarkar for their valuable assistance in establishing the CRISPR/Cas9 system in our laboratory. The authors gratefully acknowledge the financial support provided by the Department of Biotechnology and the Department of Science and Technology, Government of India.

