## Supporting Information for "Interplay of Cullin5-HECTD3 ubiquitin ligases regulates stability of CRAF mutant associated with hypertrophy"

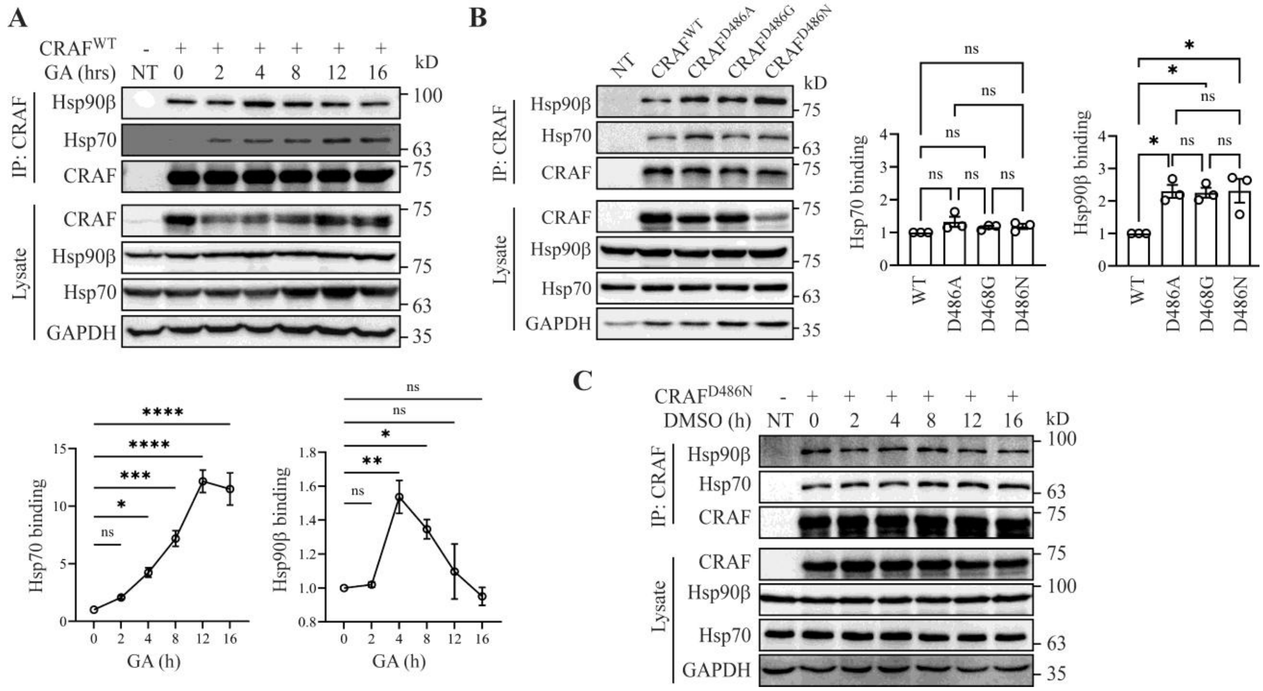


**Figure S1: Comparative analysis of chaperone interactions with WT and mutant CRAF in instability-driven dynamics. (A)** $CRAF^{WT}$ was overexpressed in *CRAF*$^{-/-}$ HEK293T cells and immunoprecipited with CRAF antibody following geldanamycin (GA) treatment at the indicated time points (0h to 16h) noted as GA (h). The association of Hsp90β and Hsp70 was evaluated by immunoblotting using their respective antibodies. Statistical analysis, one-way ANOVA, was performed using GraphPad Prism (n=3). Significance levels: ns (non-significant) $p > 0.05$, *$p < 0.05$, **$p < 0.01$, ***$p < 0.001$, ****$p < 0.0001$. **(B)** $CRAF^{WT}$, $CRAF^{D486A}$, $CRAF^{D486G}$ and $CRAF^{D486N}$ were overexpressed in *CRAF*$^{-/-}$ HEK293T cells and immunoprecipitated with CRAF antibody. The association of Hsp90β and Hsp70 was evaluated by immunoblotting using their respective antibodies. Statistical analysis, one-way ANOVA, was performed using GraphPad Prism (n=3). Significance levels: ns (non-significant) $p > 0.05$, *$p < 0.05$. **(C)** $CRAF^{D486N}$ was overexpressed in *CRAF*$^{-/-}$ HEK293T cells and immunoprecipitated with CRAF antibody at the indicated time points in the presence of DMSO. The association of Hsp90β and Hsp70 was evaluated by immunoblotting using their respective antibodies (n=1).

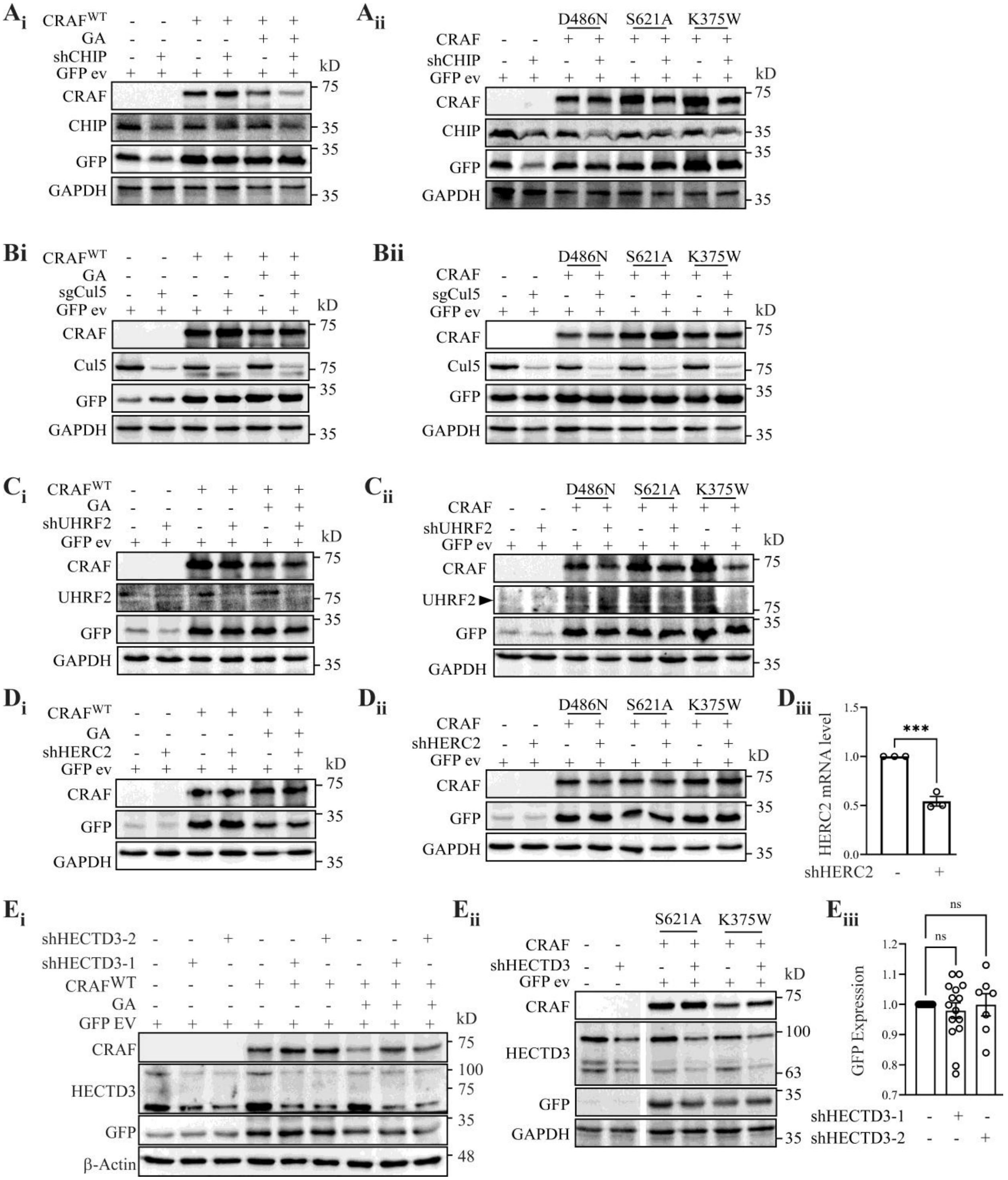


**Figure S2: Screening of ubiquitin ligases. (A-E)** Lentiviral particles encoding shRNAs targeting CHIP (A), Cul5 (B), UHRF2 (C), HERC2 (D) and HECTD3 (E) were used to infect *CRAF*$^{-/-}$ HEK293T cells. In absence and presence of GA (16h), CRAF$^{WT}$ level was assessed by using CRAF antibody compared to GFP expression which was further normalized to GFP/GAPDH ratio (Ai, Bi, Ci, Di, Ei) (n=1). Mutant CRAF (kinase-dead) expression was analyzed using an anti-CRAF antibody in cells transfected with CRAF mutants, $CRAF^{D486N}$, $CRAF^{S621A}$, and $CRAF^{K375W}$. Mutant CRAF level was assessed by using CRAF antibody compared to GFP expression which was further normalized to GFP/GAPDH ratio (Aii, Bii, Cii, Dii, Eii) (n=1). Silencing level of HERC2 was assessed by real time PCR, statistical analysis was done by students T test (n=3). Significancy level: ***$p$ < 0.001 (Diii). Level of GFP empty vector expression in shHECTD3 (1&2) condition was measured using students T test (n>3). Significancy level: ns (non-significant) $p$ > 0.05.

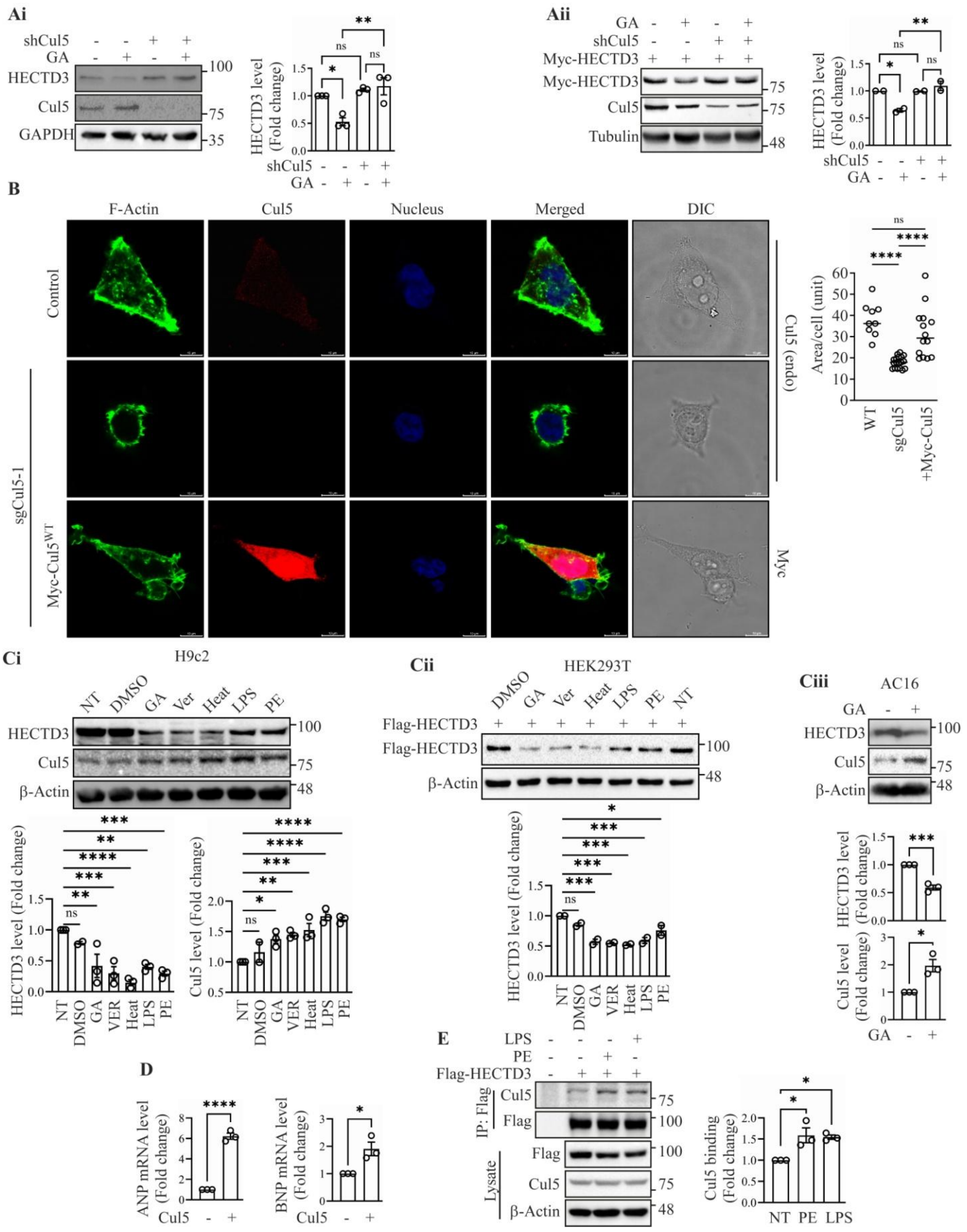

**Figure S3: Stressogenic destabilization of HECTD3 and Cul5 mediated hypertrophic induction. (A)** WT HEK293T cell and shRNA mediated Cul5-silenced HEK293T cells were treated with or without GA for 16h, and endogenous HECTD3 level was measured (Ai), similarly overexpressed HECTD3 level was determined after Myc-tagged HECTD3 was overexpressed for 48h (Aii). HECTD3 expression was assessed via western blot using anti-HECTD3 antibody (Ai) and anti-Myc antibody (Aii). Statistical analysis was performed using one-way ANOVA in GraphPad Prism for endogenous (n=3) and overexpressed (n=2) HECTD3. Significance levels: ns (non-significant) $p > 0.05$, *$p < 0.05$, **$p < 0.01$. **(B)** Representative immunofluorescence images showing the size/morphological changes. WT HEK293T and sgCul5-1 polyclonal (10-12 days after sgCul5 transfection) HEK293T cells, and Myc-Cul5$^{WT}$ transfected (for 72h) sgCul5-1 polyclonal HEK293T cells were immunostained with phalloidin (Alexa-488) for F-actin filament, DAPI for nucleus, Cul5 antibody (Alexa-594) for endogenous Cul5 and Myc antibody (Alexa-594) for overexpressed Cul5. Statistical analysis was performed by measuring surface area of cells using one-way ANOVA in GraphPad Prism (n>3). Significance levels: ns (non-significant) $p > 0.05$; ****$p < 0.0001$. **(C)** H9c2 (Ci), HEK293T (Cii) and AC16 (Ciii) cells were treated with different stressogenic factors Geldanamycin (GA) 2µM for 16h, VER-155008 (Ver) 10µM for 16h, heat stress (Heat) at 42ºC for 6h, Lipopolysaccharide (LPS) 5µg/mL for 16h, and Phenylephrine (PE) 200µM for 16h indicated in the figure. Endogenous HECTD3 and Cul5 was detected by HECTD3 and Cul5 antibody respectively in case of H9c2 (Ci) and AC16 (Ciii) and overexpressed HECTD3 was detected by Flag antibody in HEK293T (Cii) cell. Statistical significance was determined for H9c2 (n=3), HEK293T (n=2), and AC16 (n=3) by one way ANOVA using GraphPad Prism. Significance level: ns (non-significant) $p > 0.05$; *$p < 0.05$, **$p < 0.01$, ***$p < 0.001$, ****$p < 0.0001$. **(D)** AC16 cells were transfected with Myc-Cul5$^{WT}$ and the mRNA was isolated and samples were subjected to real time PCR with ANP and BNP primers taking β-Actin as control gene. Statistical significance was determined by students T test using GraphPad Prism (n=3). Significance level: *$p < 0.05$, ****$p < 0.0001$. **(E)** Association of HECTD3 and Cul5 was assessed by co-immuniprecipitation. Flag-HECTD3 was overexpressed in HEK293T cells for 48h and cells were treated with PE and LPS (conc. mentioned previously). Immunoprecipitation of the HECTD3 was performed using an anti-Flag antibody, and the presence of co-immunoprecipitated Cul5 was confirmed by immunoblotting with Cul5 antibody. Statistical analysis, one way ANOVA, was performed using GraphPad Prism (n=3). Significance level: *$p < 0.05$.

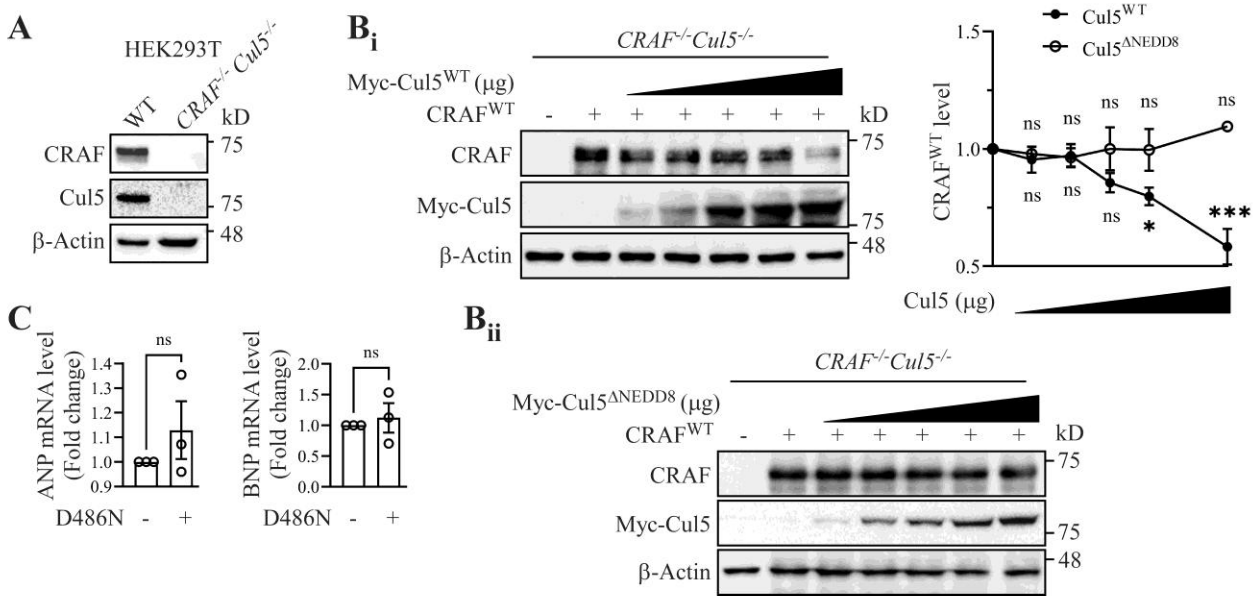


**Figure S4: Overexpression of CRAF failed to transduce hypertrophic signal. (A)** Representation of monoclonal *CRAF*$^{-/-}$*Cul5*$^{-/-}$ double-knockout HEK293T cell line, and endogenous level of the indicated proteins were checked by their respective antibodies. Here, β-Actin used as a loading control. **(B)** CRAF$^{WT}$ was overexpressed in *CRAF*$^{-/-}$*Cul5*$^{-/-}$ cells, Cul5$^{WT}$ and Cul5$^{\Delta NEDD8}$ were also overexpressed along with CRAF$^{WT}$ in a dose dependent overexpression manner (in μg) at a ratio from 0.2 to 1 (CRAF/Cul5). The level of CRAF was evaluated by immunoblotting using CRAF antibody. Statistical analysis, one-way ANOVA, was performed using GraphPad Prism (n=3). Significance levels: ns (non-significant) $p > 0.05$; *$p < 0.05$; ***$p < 0.001$; ****$p < 0.0001$. **(C)** AC16 cells were transfected with CRAF$^{D486N}$ and the mRNA was isolated and samples were subjected to real time PCR with ANP and BNP primers taking Beta actin as control gene. Statistical significance was determined by students T test using GraphPad Prism (n=3). Significance level: ns (non-significant) $p > 0.05$; *$p < 0.05$, **$p < 0.01$, ***$p < 0.001$, ****$p < 0.0001$.

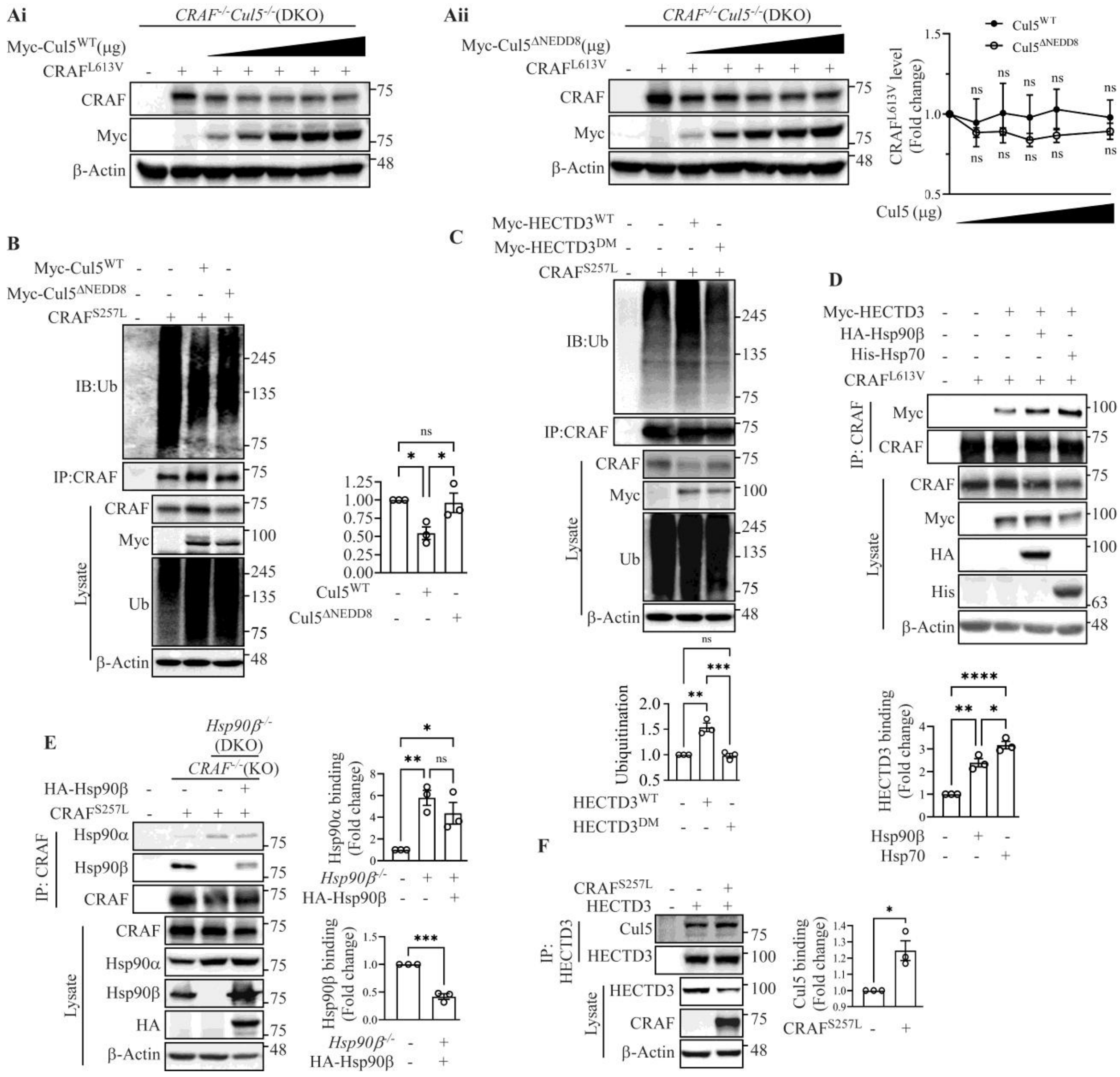


**Figure S5: Hypertrophic mutants share overlapping molecular characteristics. (A)** CRAF$^{L613V}$ was overexpressed in *CRAF*$^{-/-}$*Cul5*$^{-/-}$ cells, Cul5$^{WT}$ and Cul5$^{\Delta NEDD8}$ were also overexpressed along with CRAF$^{L613V}$ in a dose dependent overexpression manner (in µg) at a ratio from 0.2 to 1 (CRAF/Cul5). The level of CRAF was evaluated by immunoblotting using CRAF antibody. Statistical analysis, one-way ANOVA, was performed using GraphPad Prism (n=3). Significance levels: ns (non-significant) $p > 0.05$. **(B)** CRAF$^{S257L}$ was overexpressed alone or co-expressed with either wild-type Myc-Cul5 (Cul5$^{WT}$) or a neddylation-deficient mutant (Myc-Cul5$^{K724R}$/Myc-Cul5$^{\Delta NEDD8}$) followed by treatment with MG132 (5µM for 4h) in *CRAF*$^{-/-}$ *Cul5*$^{-/-}$ double knockout (DKO) cells. CRAF was immunoprecipitated using CRAF antibody and poly-ubiquitination of overexpressed CRAF was assessed. Statistical significance was determined using GraphPad Prism (n=3). Significance level: ns (non-significant) $p > 0.05$; $*p < 0.05$. **(C)** $CRAF^{S257L}$ was overexpressed in *$CRAF^{-/-}$* cells, along with Myc-$HECTD3^{WT}$ or Myc-$HECTD3^{DM}$ followed by treatment with MG132 (5μM for 4h). $CRAF^{D486N}$ was immunoprecipitated using a CRAF antibody, and poly-ubiquitination was detected by immunoblotting with a ubiquitin antibody. Statistical analysis was performed using one-way ANOVA in GraphPad Prism (n=3). Significance level: ns (non-significant) $p > 0.05$; $**p < 0.01$, $***p < 0.001$. **(D)** Association of $CRAF^{L613V}$ and HECTD3 was assessed by co-immuniprecipitation. $CRAF^{L613V}$, HA-Hsp90β, His-Hsp70 and Myc-HECTD3 were co-overexpressed in *$CRAF^{-/-}$* HEK293T cells for 48h as indicated in the figure. Immunoprecipitation of the $CRAF^{L613V}$ was performed using an anti-CRAF antibody, and the presence of co-immunoprecipitated HECTD3 was confirmed by immunoblotting with an anti-Myc antibody. Statistical analysis, one-way ANOVA, was performed using GraphPad Prism (n=3). Significance levels: $*p < 0.05$, $**p < 0.01$, $****p < 0.0001$. **(E)** Association of $CRAF^{S257L}$ and Hsp90 was assessed by co-immuniprecipitation. $CRAF^{S257L}$ was overexpressed alone in *$CRAF^{-/-}$* and *$CRAF^{-/-}$ $Hsp90\beta^{-/-}$* cell and co-overexpressed with HA-Hsp90β in *$CRAF^{-/-}Hsp90\beta^{-/-}$* for 48h as indicated in the figure. Immunoprecipitation of the $CRAF^{S257L}$ was performed using an anti-CRAF antibody, and the presence of co-immunoprecipitated Hsp90α and Hsp90β were confirmed by immunoblotting with their corresponding antibody. Statistical analysis, one-way ANOVA for Hsp90α binding and students T test for Hsp90β binding were performed using GraphPad Prism (n=3). Significance level: ns (non-significant) $p > 0.05$; $*p < 0.05$, $**p < 0.01$, $***p < 0.001$. **(F)** Association of HECTD3 and Cul5 was assessed by co-immuniprecipitation. HECTD3 and $CRAF^{S257L}$ were overexpressed in *$CRAF^{-/-}$* cells for 48h as indicated in the figure. Immunoprecipitation of the HECTD3 was performed using an anti-HECTD3 antibody, and the presence of co-immunoprecipitated Cul5 was confirmed by immunoblotting with Cul5 antibody. Statistical analysis, students T test, was performed using GraphPad Prism (n=3). Significance levels: $*p < 0.05$.

**Table of Plasmid Constructs**

| No | Construct | Information |
|---|---|---|
| 1 | pcDNA3.1-FLAG-$CRAF^{WT}$ | Gifted by Prof. Dhandapany Perundurai |
| 2 | pcDNA3.1-FLAG $CRAF^{S257L}$ | Gifted by Prof. Dhandapany Perundurai |
| 3 | pcDNA3.1-FLAG-$CRAF^{D486N}$ | Gifted by Prof. Dhandapany Perundurai |
| 4 | pcDNA3.1-FLAG-$CRAF^{L613V}$ | Gifted by Prof. Dhandapany Perundurai |

| | | |
|---|---|---|
| 5 | pcDNA3.1 FLAG-CRAF$^{S621A}$ | Generated by site directed mutagenesis |
| 6 | pcDNA3.1-FLAG-CRAF$^{K375W}$ | Generated by site directed mutagenesis |
| 7 | pcDNA3.1-FLAG-HECTD3$^{WT}$ | Generated by cloning with EcoRI and XhoI restriction enzyme from pDest40 HECTD3 vector gifted by Prof. Norbert Frey and Prof. Ashraf Yusuf Rangrez. |
| 8 | pcDNA3.1-FLAG-HECTD3$^{DM}$ | Generated by site directed mutagenesis (C744A & C823A) |
| 9 | pcDNA3.1-Myc-HECTD3$^{WT}$ | Generated by cloning with EcoRI and XhoI restriction enzyme from pDest40 HECTD3 vector gifted by Prof. Norbert Frey and Prof. Ashraf Yusuf Rangrez. |
| 10 | pcDNA3.1-Myc-HECTD3$^{DM}$ | Generated by site directed mutagenesis (C744A & C823A) |
| 11 | pcDNA3-Myc-Cul5$^{WT}$ | Addgene (#19895) |
| 12 | pcDNA3-Myc-Cul5$^{\Delta NEDD8}$ | Generated by site directed mutagenesis (K724R) |
| 13 | pcDNA3-HA-Hsp90β$^{WT}$ | Addgene (#22487) |
| 14 | pcDNA3-HA-Hsp90β$^{DN}$ | Addgene (#22480) |
| 15 | pcDNA5/FRT/TO His-HSPA1A | Addgene (#19537) |
| 16 | pLKO.1-TRC cloning vector | Addgene (#10878) |
| 17 | Scramble shRNA | Addgene (#1864) |
| 18 | psPAX2 | Addgene (#12260) |
| 19 | pMD2.G | Addgene (#12259) |
| 20 | shRNA-pLKO.1-TRC-Puro | All the plasmids of shRNA were generated by cloning with AgeI and EcoRI restriction enzyme |
| 21 | pSpCas9(BB)-2A-Puro (PX459) V2.0 | Addgene (#62988) |
| 22 | sgRNA-PX459-puro | All the plasmids of sgRNA were generated by cloning with BbsI restriction enzyme |

**List of Primers**

Table 1: shRNA primers

| No. | Primer | Sequence |
|---|---|---|
| 1 | shHECTD3-1_F | 5'CCGGGCGGGAACTAGGGTTGAATTTCTCGAGAAATTCAACCCTAGTTCCCGCTTTTTG 3' |
| 2 | shHECTD3-1_R | 5'AATTCAAAAAGCGGGAACTAGGGTTGAATTTCTCGAGAAATTCAACCCTAGTTCCCGC 3' |
| 3 | shHECTD3-2_F | 5'CCGGGCAGTCTTCACCCAGGTATATCTCGAGATATACCTGGGTGAAGACTGCTTTTTG 3' |
| 4 | shHECTD3-2_R | 5'AATTCAAAAAGCAGTCTTCACCCAGGTATATCTCGAGATATACCTGGGTGAAGACTGC 3' |
| 5 | shCul5_F | 5'CCGGGACACGACGTCTTATATTACTCGAGTAATATAAGACGTCGTGTCTTTTTG 3' |
| 6 | shCul5_R | 5'AATTCAAAAAGACACGACGTCTTATATTACTCGAGTAATATAAGACGTCGTGTC 3' |
| 7 | shHERC2_F | 5'CCGGGGAAAGCACTGGATTCGTTCTCGAGAACGAATCCAGTGCTTTCCTTTTTG 3' |
| 8 | shHERC2_R | 5'AATTCAAAAAGGAAAGCACTGGATTCGTTCTCGAGAACGAATCCAGTGCTTTCC 3' |
| 9 | shCHIP_F | 5'CCGGGGAGATGGAGAGCTATGATGACTCGAGTCATCATAGCTCTCCATCTCCTTTTTG 3' |
| 10 | shCHIP_R | 5'AATTCAAAAAGGAGATGGAGAGCTATGATGACTCGAGTCA |

| | | |
|---|---|---|
| | | TCATAGCTCTCCATCTCC 3' |
| 11 | shUHRF2_F | 5'CCGGGGTGGAATTCATGGTCGAACTCGAGTTCGACCATGAATTCCACCTTTTTG 3' |
| 12 | shUHRF2_R | 5'AATTCAAAAAGGTGGAATTCATGGTCGAACTCGAGTTCGACCATGAATTCCACC 3' |

Table 2: sgRNA primers

| No. | Primer | Sequence | Target |
|---|---|---|---|
| 1 | sgHsp90β_F | 5’ CACCGCCTGACAGACCCTTCGAAGT 3’ | exon 2 |
| 2 | sgHsp90β_R | 5’ AAACACTTCGAAGGGTCTGTCAGGC 3’ | exon 2 |
| 3 | sgCul5-1_F | 5’ CACCGGCAAAGCCGTATCATCTTGA 3’ | exon 3 |
| 4 | sgCul5-1_R | 5’ AAACTCAAGATGATACGGCTTTGCC 3’ | exon 3 |
| 5 | sgCul5-2_F | 5’ CACCGAGGCCCAGCAAAAATTCATC 3’ | exon 4 |
| 6 | sgCul5-2_R | 5’ AAACGATGAATTTTTGCTGGGCCTC 3’ | exon 4 |
| 7 | sgCRAF-1_F | 5’ CACCGACTGATGCTGCGTCTTTGAT 3’ | exon 3 |
| 8 | sgCRAF-1_R | 5’ AAACATCAAAGACGCAGCATCAGTC 3’ | exon 3 |

Table 3: Real time quantitative PCR primers

| No. | Primer | Sequence |
|---|---|---|
| 1 | HERC2_F | 5' CGGTATTTCTCTGCCTCCTG 3' |
| 2 | HERC2_R | 5' ACTCCTGCAACAGCTCACTG 3' |
| 3 | ANP_F | 5' CAGCAAGCAGTGGATTGCTCCT 3' |

| | | |
|---|---|---|
| 4 | ANP_R | 5' TCTGCGTTGGACACGGCATTGT 3' |
| 5 | BNP_F | 5' TGGAAACGTCCGGGTTACAGGA 3' |
| 6 | BNP_R | 5' TCCGGTCCATCTTCCTCCCAAA 3' |
| 7 | β-Actin_F | 5' CAAAGACCTGTACGCCAACA 3' |
| 8 | β-Actin_R | 5' TCAGGAGGAGCAATGATC 3' |

**Reagent and Chemicals**

| No. | Item | Cat no. | Company name |
|---|---|---|---|
| 1 | Geldanamycin (ant-gl-5) | | Invivogen |
| 2 | Ver-155008 | SML0271 | Sigma |
| 3 | MG132 | M7449 | Sigma |
| 4 | LPS | L4391 | Sigma |
| 5 | PMSF | P1250000 | Merck |
| 6 | Phenylephrine hydrochloride | P1250000 | Merck |
| 7 | Protease inhibitor | | Roche, Thermo |
| 8 | Phosphatase inhibitor | | Roche, Thermo |
| 9 | Ultra-Pure Tris | | MP, SRL, VWR |
| 10 | Molecular Biology grade Ethanol | | Merck, Himedia |
| 11 | Formaldehyde | | Merck, SRL |
| 12 | Vectashield | | Himedia |
| 13 | Phosphate Buffered Saline (PBS) | | Himedia |
| 14 | Dehydrated alcohol | | Himedia, SRL |
| 15 | Rectified spirit | | Bengal Chemical and Pharmaceutical Works Ltd, India |

| | | | |
|---|---|---|---|
| 16 | Tissue culture plastics | | TARSONS, BD Falcon, NUNC |
| 17 | Dimethyl Sulphoxide (DMSO) | | SRL |
| 18 | EDTA | | SRL |
| 19 | SDS | | SRL |

**Antibodies used**

| No. | Antibody name | Cat no. | Company name |
|---|---|---|---|
| 1 | Anti-FLAG | F1804 | Sigma |
| 2 | Anti-Myc | 05-724 | Merck |
| 3 | Anti-GFP | 18814460001 | Roche |
| 4 | Anti-HA | H3663 | Sigma |
| 5 | Dynabeads Protein G | 10004D | Thermo |
| 6 | Protein A Sepharose bead | GE17-0780-01 | GE Healthcare |
| 7 | Anti-CRAF | 610152 | BD Bioscience |
| 8 | Anti-β-actin | BB-AB0024 | Biobharati |
| 9 | Anti-β-actin | 15G5A11/E2 | Invitrogen |
| 10 | Anti-GAPDH | BB-AB0060 | Biobharati |
| 11 | Anti-GAPDH | MA5 15738-2 | Invitrogen |
| 12 | Anti-α-tubulin | BB-AB0118 | Biobharati |
| 13 | Anti-Ubiquitin | BB-AB0030 | Biobharati |
| 14 | Anti-Hsp90α | 8165 | CST |
| 15 | Anti-Hsp90β | 37-9400 | ThermoFisher Scientific |

| | | | |
|---|---|---|---|
| 16 | Anti-Hsp70 | ADI-SPA-811 | ENZO |
| 17 | Anti-HECTD3 | ab241413 | Abcam |
| 18 | Anti-HECTD3 | ab173122 | Abcam |
| 19 | Anti-NIRF (anti-UHRF2) | ab28673 | Abcam |
| 20 | Anti-Cul5 | NBP1-22970 | Novus |
| 21 | Anti-Cul5 | sc-373822 | SCBT |
| 22 | Anti-CHIP | N/A | Generated |
| 23 | Anti-IgG (Mouse) purified | BB-AB004 | Biobharati |
| 24 | Anti-IgG (Rabbit) purified | BB-AB003 | Biobharati |
| 25 | Goat anti-Mouse Secondary Antibody | 31430 | Invitrogen |
| 26 | Goat anti-Mouse Secondary Antibody | 115-035-003 | Jackson Biolab |
| 27 | Goat anti-Rabbit Secondary Antibody | 111-035-003 | Jackson Biolab |
| 28 | Phalloidin with Alexa-488 | A12379 | Invitrogen |
